# Genome information processing by the INO80 chromatin remodeler positions nucleosomes

**DOI:** 10.1101/2020.11.03.366690

**Authors:** Elisa Oberbeckmann, Nils Krietenstein, Vanessa Niebauer, Yingfei Wang, Kevin Schall, Manuela Moldt, Tobias Straub, Remo Rohs, Karl-Peter Hopfner, Philipp Korber, Sebastian Eustermann

**Author notes:** These authors contributed equally (Elisa Oberbeckmann, Nils Krietenstein).

## Abstract

The fundamental molecular determinants by which ATP-dependent chromatin remodelers organize nucleosomes across eukaryotic genomes remain largely elusive. Here, chromatin reconstitutions on physiological, whole-genome templates reveal how remodelers read and translate genomic information into nucleosome positions. Using the yeast genome and the multi-subunit INO80 remodeler as a paradigm, we identify DNA shape/mechanics encoded signature motifs as sufficient for nucleosome positioning and distinct from known DNA sequence preferences of histones. INO80 processes such information through an allosteric interplay between its core- and Arp8-modules that probes mechanical properties of nucleosomal and linker DNA. At promoters, INO80 integrates this readout of DNA shape/mechanics with a readout of co-evolved sequence motifs via interaction with general regulatory factors bound to these motifs. Our findings establish a molecular mechanism for robust and yet adjustable +1 nucleosome positioning and, more generally, remodelers as information processing hubs that enable active organization and allosteric regulation of the first level of chromatin.

---

The packaging of DNA with histones into nucleosomes underpins the maintenance and regulation of genome information in eukaryotes^1,2^. Genome-wide mapping of chromatin revealed highly-defined patterns of nucleosomes carrying a combinatorial landscape of histone variants and modifications^3–8^. These patterns entail well-positioned nucleosomes, which occupy the same genomic position across a cell population and even adopt equivalent positions relative to genomic sites of equivalent function like transcription start sites (TSS)^6,7^. Most prominently, nucleosome-depleted regions (NDRs) at promoters of active or poised genes are flanked by a well-positioned hallmark nucleosome (+1 nucleosome) that is the first in a regular nucleosome array over the transcribed region^9^. These stereotypic NDR-array patterns are conserved from yeast to man, and changes within their configuration play a pivotal role in transcriptional regulation, e.g., during cell differentiation and stress response^10,11^. Understanding the fundamental molecular determinants of nucleosome positioning is likely to reveal core principles by which genome regulation occurs.

A nucleosome position is defined by the DNA sequence that is wrapped around the histone octamer^12^. While this DNA sequence always answers the question “Where is this nucleosome?”, it may, but need not, answer the question “How was the nucleosome placed there?”. Histone octamers may form nucleosomes virtually at any DNA sequence position in the genome^13^. A molecular mechanism that consistently places a nucleosome at a particular genome position across a cell population must select this position against competing positions. This selection may be based on genetic information encoded within DNA sequence or on epigenetic information like histone modifications and variants or other chromatin-associated factors. Regarding DNA sequence information, pioneering studies proposed two mechanisms (Fig. 1a). One mechanism relies on the intrinsic specificity of nucleosomes to preferentially assemble on DNA sequences that favor the wrapping around the histone octamer (“genomic code for nucleosome positioning”)^14,15^. In this case, the nucleosomal DNA sequence directly determines the position. The other mechanism requires DNA sequence-specific binding of a barrier factor, to which one or several nucleosomes are aligned by statistical positioning regardless of the octamer-bound DNA sequences^16^. The principal difference between these two mechanisms illustrates two extremes, which pertain to the central question whether DNA sequence information directly or indirectly determines a nucleosome position. If directly, the nucleosome positioning mechanism reads out the DNA sequence information at the resulting nucleosome position itself. If indirectly, DNA sequence is read somewhere else, and the resulting positioning information is relayed by alignment mechanisms that position nucleosomes relative to barriers and other nucleosomes. In this case, the DNA sequence bound by the histone octamer would define, but not directly determine, the genomic position of a nucleosome.

**Figure 1.**
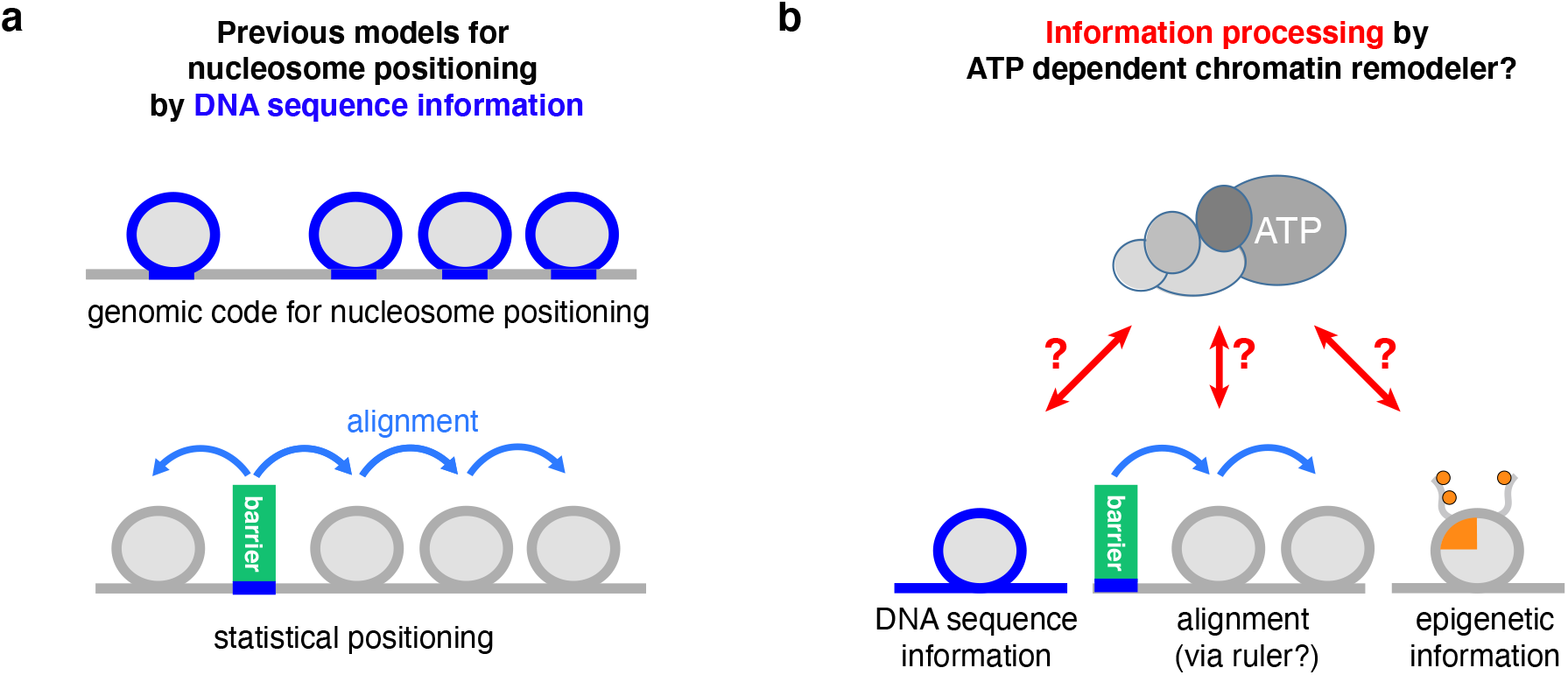
Models for nucleosome positioning mechanisms. **a** Genomic code for nucleosome positioning^14,15^ and statistical positioning^16^ are two previous models, which exemplify a direct versus indirect *role, respectively*, of DNA sequence information for determining nucleosome positioning. **b** In light of the decisive role of ATP-dependent chromatin remodelers in nucleosome positioning^24,28,29,67^, we asked if and how these large, macro-molecular machines actively process (epi)genetic information together with their own remodeler-specific information into stable nucleosome positioning.

In recent years, it has become clear that the pure versions of these two mechanistic extremes fail to explain nucleosome positioning *in vivo*. Intrinsic histone octamer preferences, as operationally assessed by salt gradient dialysis (SGD) reconstitution from purified DNA and histones^13^, cannot recapitulate NDR-array patterns *in vitro*^17,18^, and inter-nucleosomal distances (spacing) are independent of nucleosome density *in vivo*^19,20^ and *in vitro*^18,21^ in contrast to predictions of the statistical positioning mechanism^16,22^.

Instead, ATP-dependent chromatin remodelers have now been established as decisive nucleosome positioning factors by studies both *in vivo* and *in vitro*. Chromatin remodelers often form multisubunit macromolecular complexes and are grouped into four families: INO80/SWR1, SWI/SNF, ISWI, CHD. By using energy derived from ATP hydrolysis, remodelers alter histone-DNA interactions resulting in nucleosome translocation (sliding), ejection, and reconfiguration^23^. Mutations in genes encoding remodeler subunits, especially combined mutations, lead either to compromised nucleosome patterns and composition, or are lethal^20,24–28^. Complementary to genetic studies, cell-free reconstitutions provided direct evidence for the critical role of chromatin remodelers in nucleosome positioning and allowed to distinguish remodeler contributions from those of other factors, like the transcription and replication machinery^18,29^. Nucleosomes were assembled by SGD, even for an entire genome with yeast genomic DNA fragments or plasmid libraries^17,18,29,30^. The largely non-physiological nucleosome positions generated by SGD were turned in an ATP-dependent manner into *in vivo*-like NDR-array patterns either by addition of whole cell extracts^18^ or, remarkably, also by addition of remodelers purified from yeast^29^. For example, addition of yeast INO80 or SWI/SNF-type RSC remodeling complexes to SGD chromatin generated hallmark features of *in vivo*-like nucleosome organization, +1 nucleosomes and NDRs at promoters, respectively^29^. This argued for a remodeler-mediated direct readout of positioning information, possibly involving DNA sequence features^29,31^ and epigenetic information^23^. Notably, various remodelers contain reader domains of histone marks, while most of them lack classical sequence-specific DNA binding domains. This led to the proposal that remodelers, similar to histones, may recognize sequence dependent structural features of DNA such as DNA shape^32,29^. Ample and growing evidence for transcription factors underscores the functional relevance of DNA shape features in genome regulation^33^. Such features might be relevant at poly(dA:dT)-rich promoter sequences, which have been implicated in regulation of RSC activity at the NDR^31,29^, while we hypothesized that DNA shape might also play a role during +1 nucleosome positioning by INO80^29^. In contrast, other remodelers, such as the yeast ISW1a and ISW2 complexes could not generate *in vivo*-like nucleosome positions on their own but required sequence readout by other factors. So-called “general regulatory factors” (GRFs) are sequence-specific DNA binding proteins, often essential for viability and involved in transcription or replication regulation via their impact on chromatin organization^34–36^. Addition of purified GRFs, e.g., yeast Reb1 or Abf1, enabled the ISW1a and ISW2 remodelers to align regular nucleosome arrays relative to the GRF binding sites^29^. This argued in turn for remodeler-mediated readout of sequence information via processive alignment at GRFs as well as among nucleosomes, possibly involving a protein ruler^37^.

Although cell-free reconstitution and genetic studies established the critical importance of remodelers in determining the genomic organization of nucleosomes, the dissection of the underlying molecular mechanism and the required information has proven difficult. Recent structural work shed light onto the architecture of different remodelers and how they might act on mono-nucleosomes ^38^. However, there remains the conundrum that the principal remodeler activity of mobilizing nucleosomes must be regulated such that it results in stable nucleosome positions relative to genomic sequence.

In this study, we directly addressed this fundamental conundrum by asking which kind of DNA sequence, histone, barrier or other epigenetic information provides the required input, and how remodelers turn this information input into stable nucleosome positioning (Fig. 1b). We advanced whole genome reconstitutions into a fully recombinant, *de novo* approach. In this system full biochemical control is established by using recombinant components in conjunction with high resolution structural information enabling the identification of remodeling mechanisms. Not only the core mechanism of remodelers, as studied so far mainly in mono-nucleosome assays, but also the extended functions arising from remodeling of chromosomal multi-nucleosome substrates as well as the readout of physiological genomic DNA sequences and other nucleosome positioning information can be assessed at a detailed mechanistic level. We used the yeast genome and the multi-subunit structure of the INO80 complex as a paradigm to identify and probe the information and mechanism by which remodelers read information and translate it into stable nucleosome positions. In the accompanying study (Oberbeckmann & Niebauer et al.), we addressed how remodelers propagate nucleosome positioning information via an alignment mechanism to generate phased and regular nucleosomal arrays. Taken together, our data reveal that and how remodelers are information processing hubs. Genome information encoded within DNA shape/mechanics as well as in DNA sequence motifs bound by barrier factors is actively read out by the remodelers and integrated via the allosteric interplay of their molecular machinery into nucleosome positions.

## Results

### A fully recombinant approach for *de novo* whole-genome reconstitutions

To explore how ATP-dependent chromatin remodelers place nucleosomes at *in vivo*-like positions, we advanced whole-genome reconstitutions^18,29,30^ into a fully recombinant *de novo* approach (Fig. 2a). We established recombinant production of highly active and stoichiometric INO80 complex (Supplementary Fig. 1a,b) and performed whole-genome reconstitutions using recombinant histones and a fully-defined yeast genomic plasmid library^39^. This leverages, compared to previously used ill-defined plasmid libraries, endogenous fly embryo histones and endogenous purifications of remodelers^29^, the full potential of biochemical systems: (1) A fully defined 15-subunit *S. cerevisae* INO80 complex, amendable for structure-guided mutagenesis, (2) histones without posttranslational modifications (PTMs) and amendable for mutagenesis, and (3) defined DNA templates for chromatin assembly. We used MNase-seq to measure resulting nucleosome positions.

**Figure 2.**
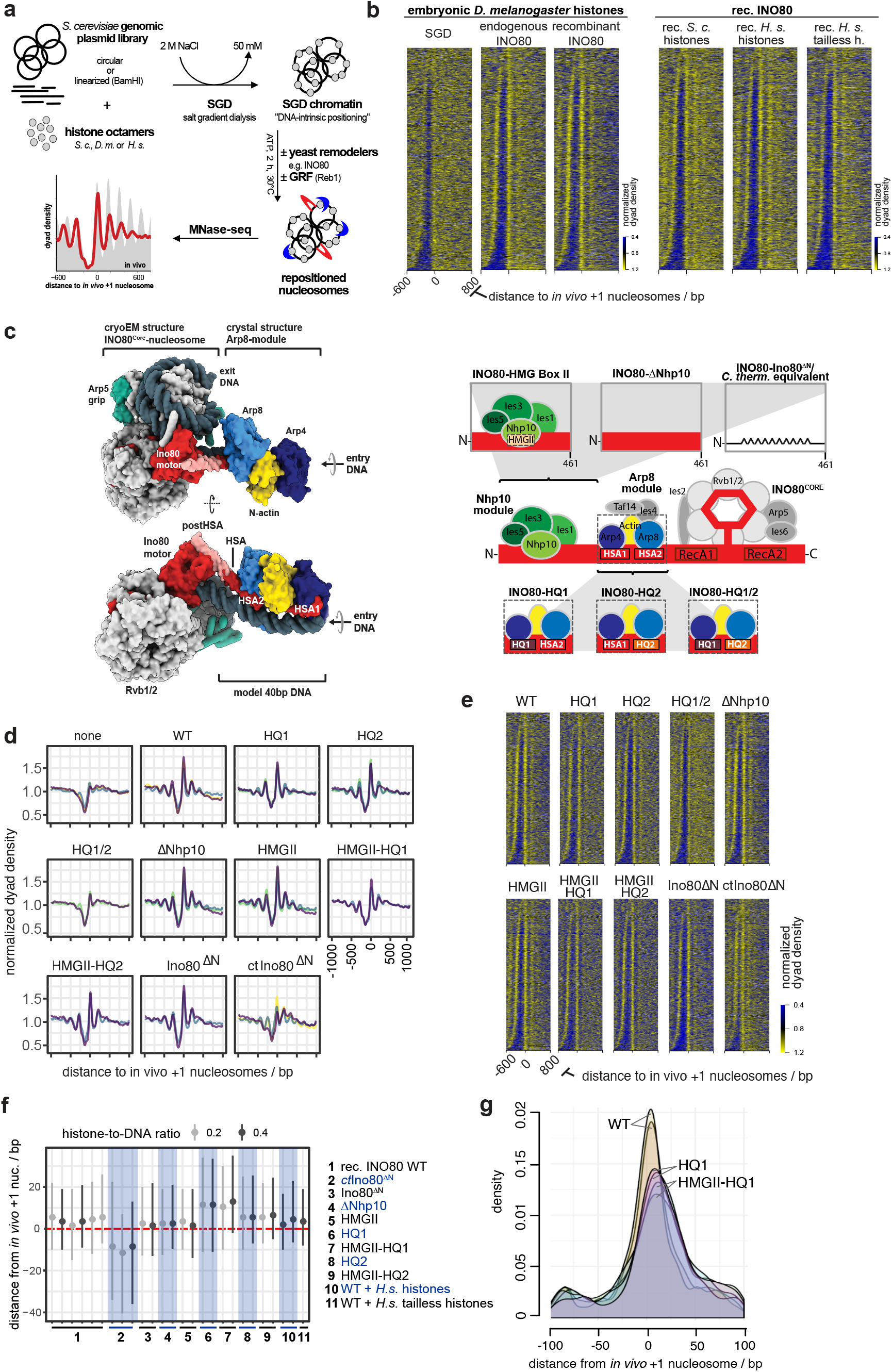
Fully recombinant genome-wide reconstitution of nucleosome positioning by INO80. **a** Overview of genome-wide *in vitro* chromatin reconstitution system. **b** Heat maps of MNase-seq data for SGD chromatin assembled with embryonic or recombinant (rec.) histones from the indicated species (“*H. s.*” abbreviates *Homo sapiens*, “*S. c.*” abbreviates *Saccharomyces cerevisiae*.) and remodeled with endogenous or recombinant *S. cerevisiae* INO80 complex as indicated. Heat maps are aligned at *in vivo* +1 nucleosome positions and sorted by NDR length. Single replicates are plotted, see Supplementary Figure 1c for all replicates. **c** Left panel: Composite model of INO80 based on high resolution cryoEM structure of ctINO80 core in complex with a mono-nucleosome^43^ and X-ray structure of Arp8 module modeled on 40bp linker DNA^46^. Images taken from Knoll et al. ^46^. Right panel: Schematic of INO80 complex submodule and subunit organization (middle) with zoom into Nhp10 (top) or Arp8 module (bottom) showing three mutant versions each. **d** Composite plots of MNase-seq data of individual replicates for SGD chromatin incubated with the indicated recombinant WT (WT) or mutant INO80 complexes (as in panel c) from *S. cerevisiae* or *C. thermophilum* (*ct*INO80^ΔN^). **e** Heat maps of MNase-seq data for samples as in panel d. **f** Distributions of distances between +1 nucleosome positions determined by paired-end sequencing after reconstitution by the indicated combinations of INO80 complexes and histones at the indicated histone-to-DNA mass ratio relative to *in vivo* +1 nucleosome positions. Dots mark the medians, vertical lines the interquartile distances. Alternating white and grey vertical zones group replicates of the indicated remodeler/histone combinations. **g** Density distributions of MNase-seq reads relative to *in vivo* +1 nucleosome positions of samples with INO80 WT, HQ1 and HMGII-HQ1 mutant complexes as in panel f.

### DNA sequence and globular histone octamer information is sufficient for *in vivo*-like +1 nucleosome positioning by INO80

This recombinant system enabled us to identify the minimal information for nucleosome positioning by INO80. Consistent with its localization and function *in vivo*^40^, INO80 positions *in vivo*-like +1 nucleosomes adjacent to NDRs (Fig. 2b,^29^). As equally pronounced +1 nucleosome positioning activity was observed for recombinant as for endogenous INO80 (Fig. 2b, left), we concluded that no yeast-specific PTMs were required and no co-purified yeast contaminant was responsible. To control the specificity of the highly pure INO80 complex (Supplementary Fig. 1a,b), we assayed an INO80 complex which carries a Walker B motif mutation within its Ino80 ATPase motor protein (Supplementary Fig. 1c) and excluded that nucleosome positioning activity was due to any co-purifying factor(s) from insect cells. Intriguingly, our recombinant whole-genome reconstitutions established conditions, under which INO80 generated extensive nucleosome arrays (e.g., upon addition of Reb1, see below). This served as starting point for the study of nucleosome spacing mechanisms (accompanying paper by Oberbeckmann & Niebauer et al.).

Next, we asked whether epigenetic information derived from histone modifications or variants was required for +1 nucleosome positioning. Histone variants, for example H2A.Z, may alter direct, sequence-dependent interactions of the histone octamer^41^. However, compared to SGD chromatin prepared with endogenous fly histones, using either recombinant human or yeast histones resulted in very similar nucleosome positioning by INO80 (Fig. 2b, right). Patterns were less pronounced with yeast histones, which we attributed to their known propensity to form less-stable nucleosomes^42^. As the species-origin of the histones did not matter much, we went more minimalistic and asked if just the globular histone domains were sufficient. SGD chromatin with recombinant tailless human histones still allowed INO80 to position *in vivo*-like +1 nucleosome position (Fig. 2b, right). We observed increased sliding rates with tailless compared to full-length histone nucleosomes (Supplementary Fig. 1d) consistent with previous studies^43–45^. Nonetheless, this increased sliding rate did not abrogate formation of the steady state nucleosome positioning pattern.

Taken together, we concluded that neither histone modifications nor histone variants nor histone tails nor yeast-specific modifications are absolutely required for INO80 principal activity to position in vivo-like +1 nucleosome. Consequently, INO80 can generate such positioning solely by processing information from genomic DNA sequences and the globular histone octamer. Nonetheless, a readout of epigenetic information by remodelers is expected to play a pivotal role in the regulation of nucleosome positioning, e.g., in response to changes in the cellular environment, as discussed further below.

### Structure-based site-directed mutagenesis probes nucleosome positioning by INO80

Having identified a minimal set of components, from which INO80 derives nucleosome positioning information, we set out to specify this information and to dissect the molecular mechanism, by which it was processed. To this end, we leveraged high-resolution structures of INO80^43,45,46^ and asked which remodeler elements might function as reader of genome information.

Recent structural and biochemical studies revealed an extended configuration of the INO80 multi-subunit architecture on mono-nucleosomes (Supplementary Fig. 1f): the INO80 core module (Ino80 protein containing the Snf2-type ATPase, Ies2, Ies6, Arp5, Rvb1, Rvb2) engages the nucleosome core particle^43,45^, the nuclear-actin containing Arp8 module (Ino80-HSA domain, Arp8, Arp4, nuclear actin, Ies4 and Taf14) binds along 40-50 bp of linker DNA at the entry site^43,45,47^, while the species-specific Nhp10 module (Nhp10, Ies1-3 and Ies5) bound to the Ino80 N-terminal region is located at the distal site of INO80’s linker DNA footprint^47^. Linker DNA binding by the Arp8 and Nhp10 modules was proposed to provide a DNA linker length dependent sensor that is allosterically coupled to processive nucleosome translocation catalyzed by the INO80 core^46–48^. *In vivo* ChIP-exo mapping suggested a highly similar INO80 configuration at +1 nucleosomes with the Arp8 or Nhp10 modules located at adjacent promoter regions^40^. Thus, we reasoned that these INO80 modules are prime candidates for reading genomic DNA sequence information.

To test this hypothesis, we targeted candidate INO80-DNA interactions based on the high-resolution cryoEM and X-ray structures of the INO80 core and Arp8 module, respectively, as well as on homology modeling of the structurally less well characterized Nhp10 module. For the INO80 core, we tested the role of ATP hydrolysis by the hetero-hexameric AAA^+^-ATPase Rvb1/2 (Fig. 2c, Supplementary Fig. 1c), which structurally organizes the nucleosome core binding and remodeling unit of INO80^43,45^. For the Arp8-module, we employed the Ino80-HSA helix mutants, which contain substitutions of highly conserved lysine/arginine to glutamine residues in the HSAα1 and/or HSAα2 helices (HQ1, HQ2 and combined HQ1/2 mutants, respectively) that are important for linker DNA binding^46^ (Fig. 2c, Supplementary Fig. 1e). For the Nhp10 module, we either mutated site-specifically the HMG box II in Nhp10 based on well-known DNA binding activity of HMG box proteins or removed the entire Nhp10 module by deleting Nhp10 or truncating Ino80’s N-terminal 1-461 residues, to which this module binds (Fig. 2c, Supplementary Fig. 1e,g,h). This latter mutant corresponded to the *Chaetomium thermophilium* INO80 core complex used in the cryoEM structure^43^, which we also employed here. Nhp10 module HMGII box and Arp8-module HQ1 or HQ2 mutations were also combined (HMGII-HQ1, HMGII-HQ2 mutants, respectively) (Fig. 2c, Supplementary Fig. 1e).

### The INO80 Arp8 module is a reader of genomic sequence information

Comparison of nucleosome patterns in aligned heat map or composite plots suggested that most INO80 mutant complexes generated similar +1 nucleosome positioning as WT INO80 (Fig. 2d,e, Supplementary Fig. 1c). Rvb1/2 ATPase activity was not required (Supplementary Fig. 1c), consistent with the likely role of Rvb1/2 during INO80 biogenesis^49^. Even the heterologous *C. thermophilum* INO80 core complex (ctINO80^ΔN^) appeared to generate +1 nucleosomes on the *S. cerevisiae* genome to a remarkable extent, suggesting a conserved readout mechanism (Fig 2d,e). Only the HQ1/2 double mutant complex was substantially impaired in +1 nucleosome positioning (Fig. 2d,e), consistent with its impaired nucleosome sliding and decoupled ATPase activity^46^. The apparent robustness of INO80 +1 nucleosome positioning activity was in contrast to the nucleosome spacing activity, which was affected for most of these INO80 mutants (accompanying paper by Oberbeckmann & Niebauer et al.).

Quantification of distances between +1 nucleosome positions reconstituted *in vitro* and observed *in vivo* revealed a distinct impact of INO80 mutations (Fig. 2f,g). Paired-end sequencing enabled accurate determination of nucleosome dyad positions on individual DNA molecules, while we included also a lower histone-to-DNA mass ratio (~0.2, accompanying paper by Oberbeckmann & Niebauer et al.) than mostly used in this study (~0.4) to further reduce possible next-neighbor nucleosome effects. WT INO80 and Nhp10 module mutants generated *in vivo*-like +1 nucleosomes with remarkable precision (Fig. 2f,g), whereas INO80 complexes bearing the HQ1 mutation and the ctINO80^ΔN^ complex generated +1 nucleosome positions that deviated more from the *in vivo* positions than those generated by the other complexes (Fig. 2f). Compared to WT INO80, +1 nucleosome positioning by complexes with the HQ1 mutation was shifted by 10 bp downstream and reduced positioning precision was reflected in broadened distributions, which suggests that DNA sequences underlying *in vivo* +1 nucleosome positions correspond more to the DNA sequence preferences for nucleosome positioning of the WT versus the mutant INO80 complexes (see below). (Fig. 2f,g). Such downstream shifts, observed here for individual INO80 point mutations, were reminiscent of similar effects resulting from INO80 depletion in the context of the interplay with other remodelers *in vivo*^20,28,40,50^. Taken together, our mutational analysis of candidate DNA contacts indicated robust processing of genomic sequence information by INO80 with a decisive role of the Arp8, but not the Nhp10 module, as direct reader of genome information at promoters.

### DNA shape/mechanics readout underlies nucleosome positioning by INO80

Based on our mutational analysis, we sought to identify genomic DNA sequence features that provide positioning information. Previously, we proposed that *S. cerevisiae* INO80 might read DNA shape features of nucleosomal DNA^29^. However, this hypothesis was based on correlation and the approach limited further interpretation, mainly because we used gene ranking by MNase-seq signal strength at pre-defined +1 to +3 nucleosome regions before and after remodeling as the discriminating category. This may introduce a bias towards the starting conditions, i.e. DNA sequence preferences of histones and variations in SGD assembly conditions. Moreover, the analysis was limited to pre-defined regions and numerous other DNA sequence motifs present at gene starts, e.g., evolved in the context of transcription regulation, may have convoluted the search for positioning information.

Here, we overcame these limitations and searched for the DNA sequence features of nucleosome positioning preferences by INO80 more globally, not only at promoters, and explored by a structure-based mutational analysis the direct and causal impact of altered INO80-DNA contacts on these preferences. We established a sensitive and unbiased Principal Component Analysis (PCA)/clustering approach solely on the basis of *de novo* generated nucleosome dyad positions determined by paired-end sequencing. This enabled unsupervised PCA/clustering of a large number of datasets (e.g. replicates, different assembly degrees, various INO80 WT and mutant complexes etc.) without prior assumptions (Fig. 3a).

**Figure 3.**
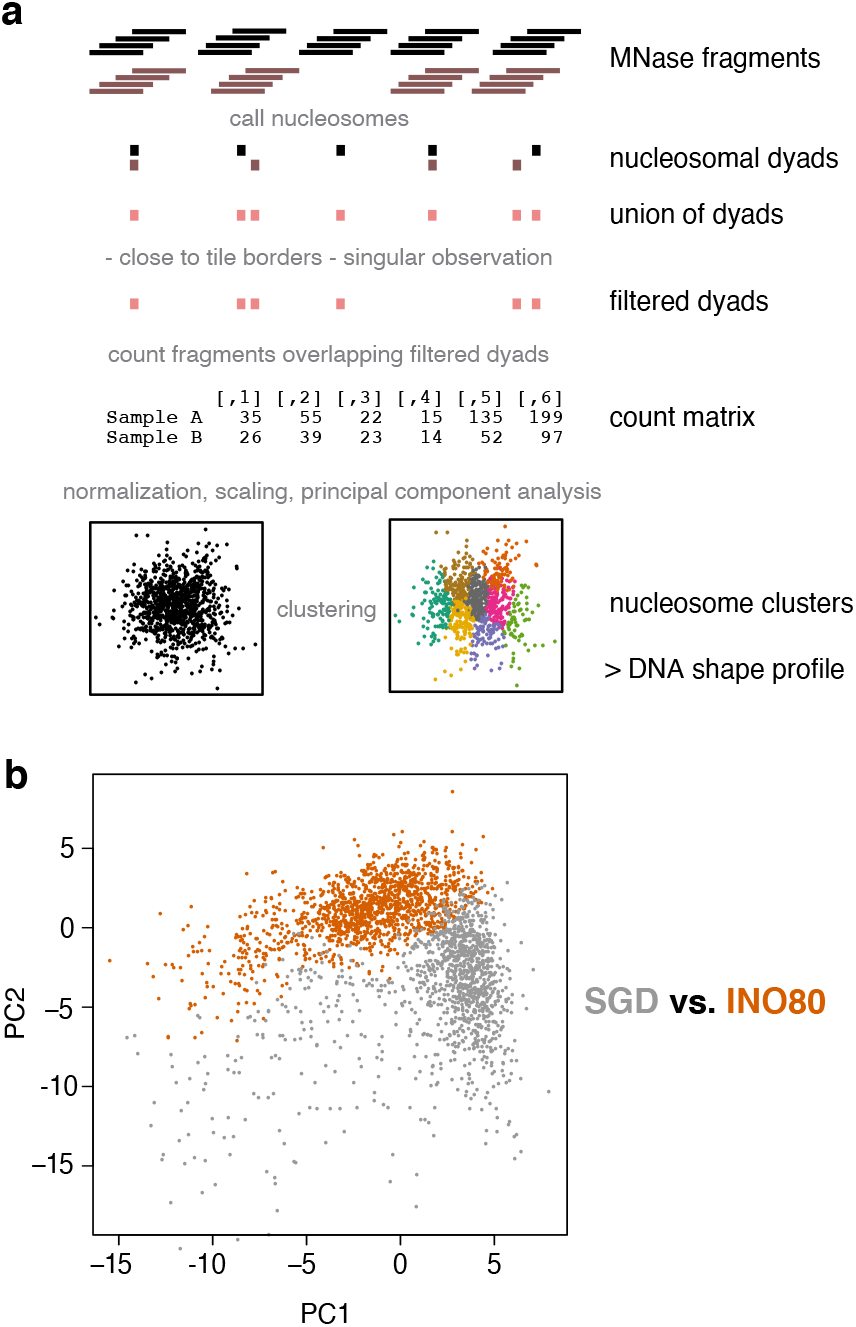
Principal Component Analysis (PCA)/clustering approach. **a** Schematic of the analysis by using two conditions (black and grey) as an example. For details see main text and Materials and Methods section. **b** Visualization of nucleosome clusters according to Principal Components 1 and 2 (PC1, PC2) for SGD chromatin (SGD) prepared with embryonic *D. melanogaster* histones at histone-to-DNA mass ratio of 0.4 alone (SGD) or after incubation with *S. cerevisiae* WT INO80 complex (INO80). INO80 remodeling alters almost the entire landscape of nucleosome positions.

Nucleosomes remodeled by WT INO80 clearly clustered differently in PCA than those assembled during SGD without remodeling (Fig. 3b), i.e. this approach could clearly distinguish positioning preferences under different conditions. The DNA sequences of different clusters did not differ in terms of sequence motifs assessed by motif search algorithms like Homer (data not shown) in contrast to previous studies of an isolated, truncated construct of human INO80 HSA domain that indicated sequence-specific DNA binding^51^.

However, DNA sequence information need not result in classical sequence motifs but may correspond to DNA shape features that are encoded in a more redundant way, i.e., rather disparate sequences may share similar shape features^52^. A composite plot of the DNA shape feature propeller twist of SGD-reconstituted versus INO80-remodeled nucleosomes revealed symmetrical but strikingly different profiles (Fig. 4a), revealing distinct DNA sequence requirements for INO80- and SGD-mediated positioning. Whereas propeller twist is largely affected by the number of intra-bp hydrogen bonds, other shape features gave corresponding results (Supplementary Fig. 2a). These other shape features take into account interactions either between adjacent bp (helix twist and roll) or with additional nucleotides (minor groove width). The profile symmetry validated the shape information content as no nucleosome orientation was to be expected and symmetrical shape profiles are unlikely to occur by chance if no underlying shape feature were involved. Importantly, similar but asymmetrically distorted shape profiles were seen for nucleosomes reconstituted at positions close to *in vivo* +1 nucleosome positions and oriented relative to the direction of transcription (Fig. 4c). This shows that such pronounced DNA shape signals are also present in +1 nucleosome regions at gene promoters and strongly suggested that we identified the DNA-encoded signal for INO80-mediated +1 nucleosome positioning. The structural readout of DNA features, both in the gene promoter as well as in +1 nucleosome, is also consistent with *in vivo* binding of INO80 subunits to such regions, as observed by ChIPexo mapping^40^.

**Figure 4.**
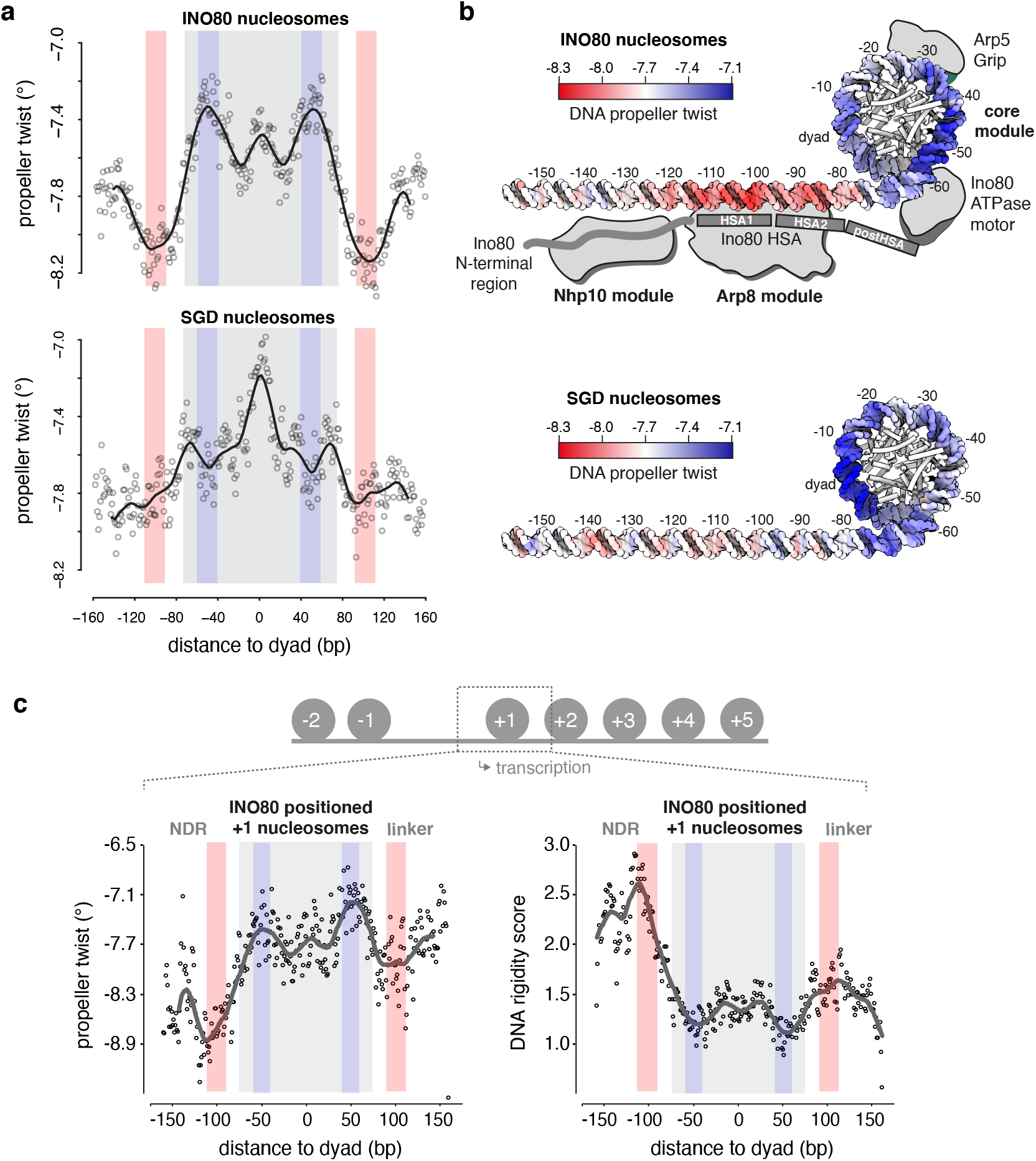
DNA shape readout underlies nucleosome positioning by INO80 and SGD. **a** Propeller twist DNA shape profiles for nucleosomal sequences of SGD chromatin with (INO80 nucleosomes) or without (SGD nucleosomes) remodeling by recombinant *S. cerevisiae* WT INO80 complex. Light red and light blue background indicate regions of major differences between SGD and INO80 profiles. Light grey background marks the location of the nucleosome core particle. **b** Red-white-blue color gradient mapping of propeller twist DNA shape profile from panel (a) on model of linker and nucleosomal core DNA. Binding architecture of INO80 is shown schematically and based on structural data^43,46^ and biochemical mapping^47^. **c** Propeller twist DNA shape and DNA rigidity profiles for INO80 positioned *+*1 nucleosomes, all with the same orientation relative to the direction of transcription. See main text and Materials and Methods for a description of the DNA rigidity score. Note that the promoter NDR around −100 bp corresponds to a rigid DNA motif, while the score indicates an increased flexibility around −55 bp between the ATPase-motor and the Arp5-grip of INO80 (see panel b).

DNA shape profiles establish a new kind of nucleosome positioning information that is distinct from previously known DNA sequence preferences of histones. The relevance of DNA shape for remodeler-mediated nucleosome positioning was further underscored by a striking congruency between our PCA/clustering data and high-resolution structural information as well as *in vivo*-mapping of INO80 subunits at gene promoters. The remodeled nucleosomes differed mostly in the ± 55 bp and ± 100 bp regions relative to the dyad (color shaded areas in Fig. 4a) where functionally important interactions with the INO80 complex are suggested by the biochemical and structural information available from INO80 in complex with mono-nucleosomes (Fig. 4b). The HSA helix at the Ino80 N-terminus contacts linker DNA at about −100 bp from the dyad^46,47^. The −55 bp region from the dyad lies between the Ino80 ATPase domain and the DNA contact point of Arp5. Both of these regions are critically important for nucleosome translocation. DNA strain build-up in the −55 bp region by successive rounds of DNA pumping by Ino80 ATPase motor is a central element of the proposed core mechanism of nucleosome translocation by INO80, while sensing of linker DNA by the Arp8 module ensures allosteric coupling of ATP hydrolysis to DNA translocation, which has been proposed to prevent back-slippage during DNA strain build up^43,47^.

This congruency immediately suggests a molecular mechanism by which an active readout not only through recognition of ground-state average DNA shape features, but also via ATP-hydrolysis driven perturbation of mechanical properties of DNA leads to the positioning of nucleosomes. The most immediate mechanical property of the double-helix is conformational flexibility. To assess this property on a genomic scale, we introduced a rigidity score that characterizes how rigid/flexible DNA is within a local region at bp resolution^33^. We considered A-tracts of consecutive ApA (TpT) or ApT bp steps as dominant factor in increasing rigidity due to strong stacking interactions combined with inter-bp hydrogen bonds in the major groove^32,53^. The rigidity score accounts for the length of A-tracts as longer runs of ApA (TpT) and ApT steps without TpA steps or G/C bp increase rigidity of a DNA fragment. We observed that DNA rigidity is correlated with DNA shape features, and the correlation remains at a consistent level across INO80 positioned nucleosomes (Supplementary Fig. 2a,b,c). This analysis reveals that +1 nucleosome positioning by INO80 involves placement of nucleosomes where DNA flexibility is increased at the −55 bp region between the ATPase motor and the Arp5 grip, while the promoter NDR region harbors a rigid DNA element where the Arp8-module is located (Fig. 4c). Intriguingly, a similarly rigid promoter DNA motif at the same distance in respect to the +1 nucleosome was also identified in a parallel study, where DNA mechanics were measured experimentally on a genomic scale via library-based DNA circularization assays^54^.

### Altered Ino80-HSA-helix-DNA contacts affect DNA shape/mechanics readout by INO80

To establish causality, we probed whether the INO80-DNA contacts and different histones would affect the readout of DNA shape/mechanics. Nucleosomes positioned by WT INO80 clustered together with those positioned by mutant complexes where mutations affected the Nhp10 module, i.e., the Ino80 N-terminus or Nhp10 module subunits including the Nhp10 HMG Box (Fig. 5a). This corroborated our results regarding nucleosome positioning in promoter regions (Fig. 2d-f) and ruled out a major role for the Nhp10 HMG box in DNA shape/mechanics readout by INO80. In contrast, all mutant complexes impaired in HSA helix-DNA contacts, either the HQ1 or HQ2 mutation and each also in combination with the HMGII mutations, generated distinct clusters of nucleosome positions (Fig. 5a). Overall shape/mechanics preferences were not much affected if endogenous fly versus recombinant human histones were used (Fig. 5b). This validated our use of fly histones for the comparisons among WT and mutant INO80 complexes in this approach.

**Figure 5.**
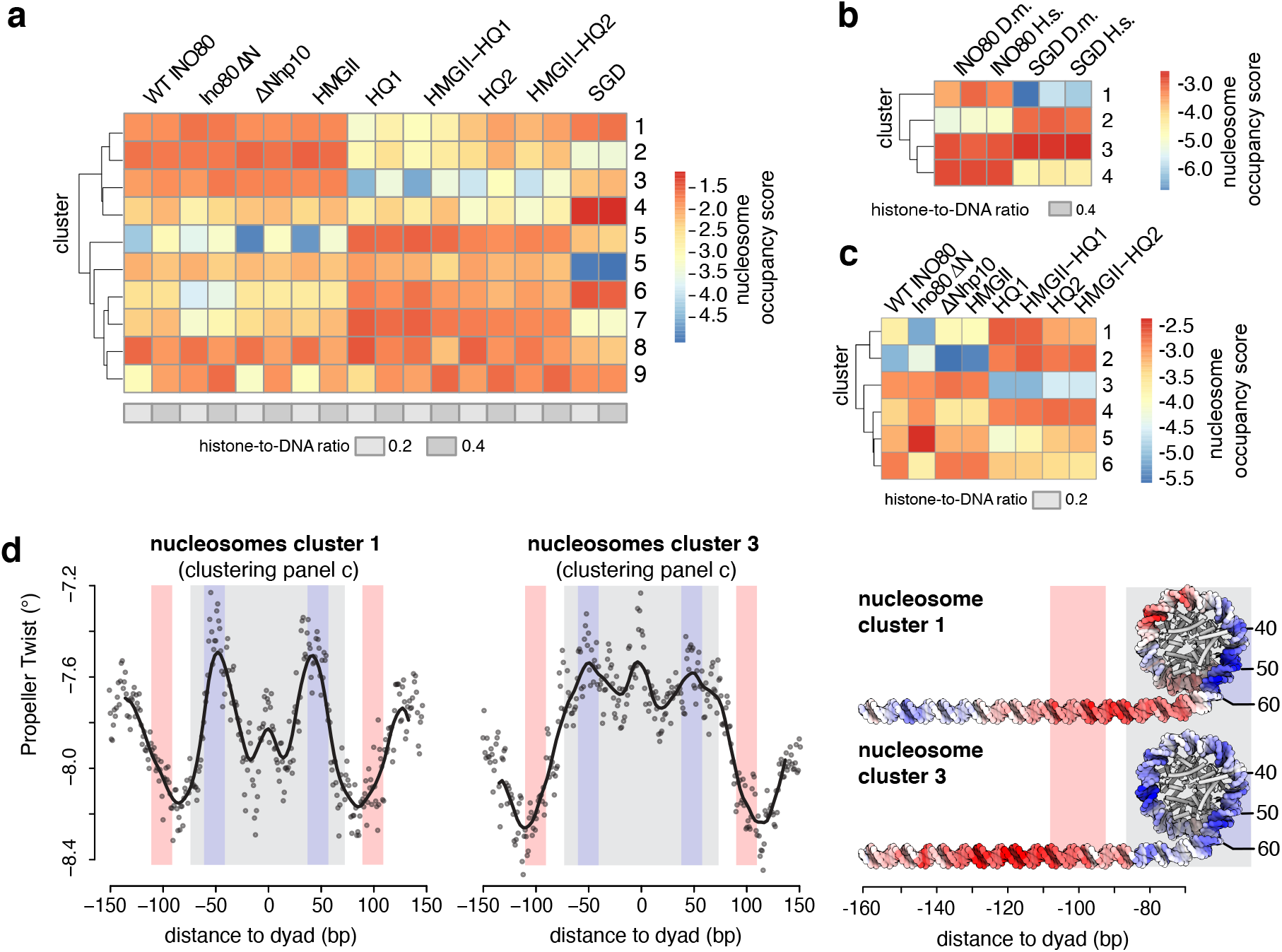
Structure-based mutations probe the DNA shape/mechanics readout by INO80. **a** Nucleosome position clusters derived from principal component analysis (PCA) of nucleosome positions of SGD chromatin with embryonic *D. melanogaster* histones at the indicated histone-to-DNA mass ratio without (SGD) or after remodeling by the indicated recombinant *S. cerevisiae* WT and mutant INO80 complexes (as in Figure 3d,e) **b** As panel a but for SGD chromatin with embryonic *D. melanogaster* (*D. m*.) vs. recombinant *H. sapiens* (*H. s*.) histones at the indicated histone-to-DNA mass ratio without (SGD) or with remodeling by recombinant *S. cerevisiae* WT INO80 complex (INO80). **c** As panel b but only with the indicated subset of samples. **d** As panel a but only for nucleosomes from the indicated clusters of panel c. Propeller twist DNA shape data mapped onto model of linker and nucleosomal DNA by using red-white-blue color gradient. See Supplementary Figure 3 for other clusters.

In total, there were three major classes of nucleosome positions, those generated by i) SGD, ii) WT INO80/Nhp10 module mutant complexes or iii) HSA helix mutant complexes (Fig. 5a). To investigate the differences in DNA sequence preferences only between the INO80 complexes and at minimal contribution of neighboring nucleosomes, we clustered only the respective samples with low assembly degree SGD chromatin (Fig. 5c) and compared the resulting DNA shape/mechanics profiles of clusters with clearly different occupancies among the INO80 complexes, e.g., cluster 1 versus 3 (Fig. 5c, Supplementary Fig. 3). Propeller twist signal profiles clearly differed between clusters that contained nucleosome positions preferentially generated by the HSA helix-mutated INO80 versus WT or Nhp10 module mutated complexes. In particular, the ± 100 bp region of the linker DNA showed a distinct shift of the propeller twist signal by more than 20 bp between cluster 1 and 3 (Fig. 5d). As this is the region where the Ino80 HSA domain contacts DNA (Fig. 4b), these data directly showed that these HSA helix-DNA contacts contributed to the DNA shape/mechanics readout during nucleosome positioning. Moreover, additional changes of propeller twist signals within the nucleosomal DNA region provided, in context of Ino80 HSA mutations, evidence for the allosteric interplay between the Arp8- and the core module of INO80^46,47^. We conclude that INO80 positions nucleosomes via a readout of DNA shape/mechanics profiles. This information and its readout are distinct from known DNA sequence preferences of histones suggesting that remodelers play an active role in translating genomic information into nucleosome positions, i.e., determine nucleosome positions through their specific molecular mechanism of remodeling.

### The DNA sequence-specific barrier Reb1 regulates nucleosome positioning by INO80

Having established that INO80 reads DNA shape/mechanics features and translates this information via specific modules into nucleosome positions, we asked next whether INO80 is also capable of processing nucleosome positioning information from DNA sequence-specific barrier factors (Fig. 1b). Reb1 is a GRF important for promoter nucleosome organization *in vivo*^26^. Sequence-specific GRFs serve, via an unknown mechanism, as nucleosome positioning alignment point for remodelers like ISW1a or ISW2^29^.

To directly test whether Reb1 binding at cognate promoter sites controls +1 nucleosome positioning by INO80, we turned again to whole-genome reconstitutions. Increasing Reb1 concentrations clearly improved nucleosome positioning by INO80 at promoters with Reb1 sites in terms of +1 nucleosome occupancy (peak height), but also in array extent and NDR depth (Fig. 6a,b, Supplementary Fig. 4a). This Reb1 effect was again independent of the histone octamer species-origin (Supplementary Fig. 4b). Detailed quantification of nucleosome spacing and array phasing at Reb1 sites and at different nucleosome densities was studied in the accompanying paper (Oberbeckmann & Niebauer et al.). *In vivo* mapping of INO80 subunits by ChIPexo^40^ indicated that INO80 adopts an extended conformation, which might bridge Reb1 binding sites and +1 nucleosomes.

**Figure 6.**
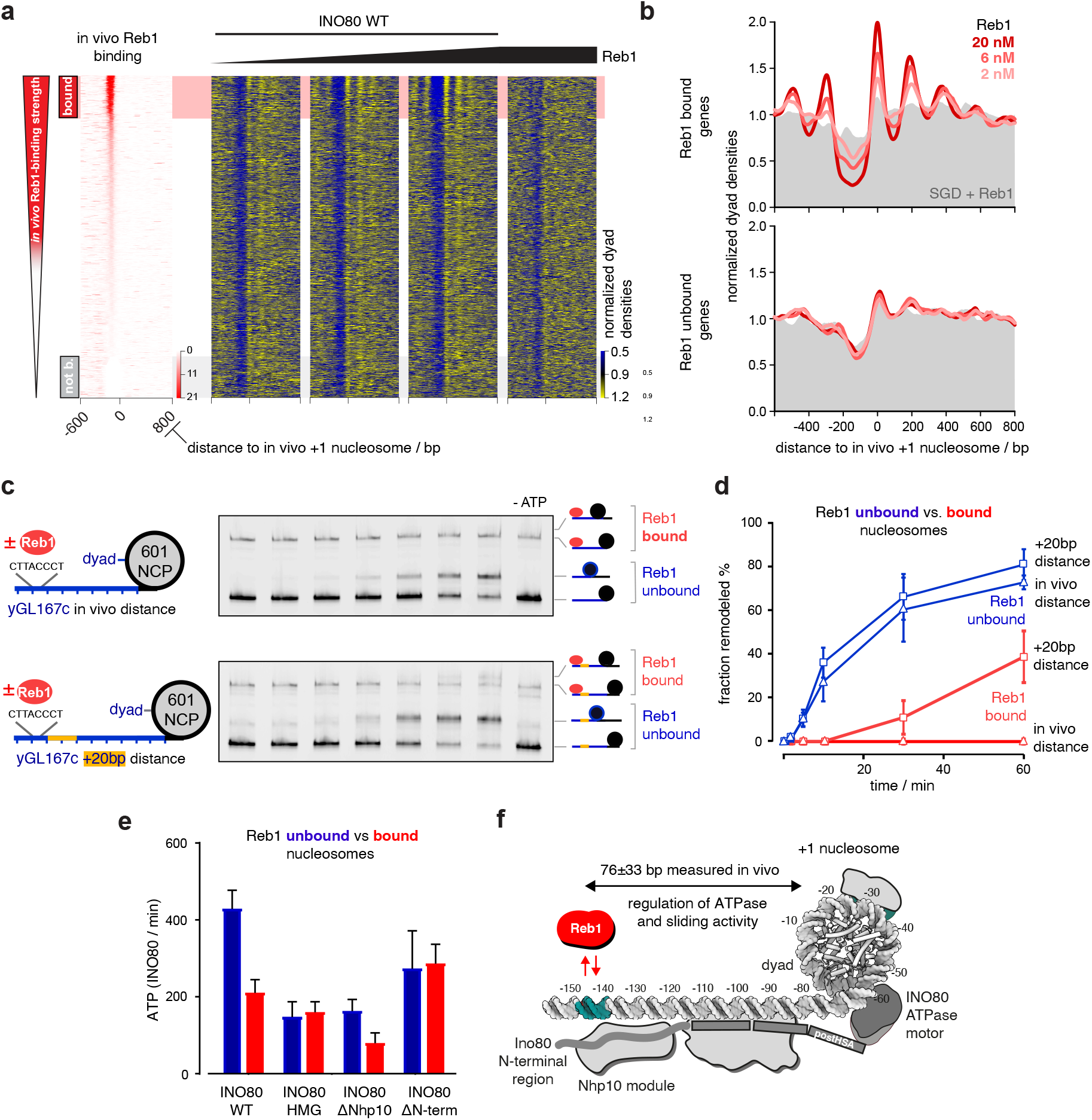
Reb1 regulates nucleosome positioning by INO80 and INO80’s ATPase and sliding activity. **a** Heat maps of MNase-seq data for SGD chromatin assembled with recombinant *H. sapiens* histones at histone-to-DNA mass ratio 0.4, incubated with recombinant *S. cerevisiae* WT INO80 and increasing concentrations of recombinant Reb1 (ramp denotes 2, 6 and 20 nM Reb1). Right most panel shows sample prepared with embryonic *D. melanogaster* histones. Heat maps are aligned at *in vivo* +1 nucleosome positions and sorted according to decreasing (top to bottom) anti-Reb1 SLIM-ChIP score (*in vivo* Reb1 binding^68^) shown in leftmost heat map. Horizontal red or grey shading highlights genes with strong or weak *in vivo* Reb1 promotor-binding, respectively. Single replicates were plotted, see Supplementary Figure 3a for all replicates. **b** Composite plots of MNase-seq data as in panel A averaged over genes highlighted in red (top) or grey (bottom) in panel (a). **c** Left: mononucleosome substrate design with 80 bp (top) or 100 bp DNA overhang (bottom) taken from a promoter (yGL167c) with clear +1 nucleosome positioning by just INO80 *in vitro* and INO80 bound *in vivo* ^40^. Guided by its dyad positions, we replaced the genomic +1 nucleosome sequence of yGL167c with a 601-nucleosome positioning sequence. Right: Native PAGE nucleosome sliding assay for indicated mononucleosome and Reb1 concentrations, and 10 nM recombinant *S. cerevisiae* WT INO80 for yGL167c-NCP601 (top) or yGL167c-20-NCP601 (bottom). “-ATP” denotes 60 min time point without ATP. **d** Quantification of sliding assays from the middle panel and two other replicates. Traces in red show data in the presence of Reb1. Error bar shows SD between replicates. **e** NADH-based ATPase assay for the 25 nM mononucleosomes and 10 nM recombinant *S. cerevisiae* WT and mutant INO80 complexes alone or with Reb1 at equimolar ratio to mononucleosome respectively. **f** Structural data^43,46^ and biochemical mapping^47^ suggest a putative binding architecture of INO80 which might bridge Reb1 and +1 nucleosomes. Allosteric communication occurs across a distance of more than 70 bp (median of 76±33bp *in vivo* measured by SLIM-ChIP^69^ and MNase-seq).

To directly address whether INO80 relays positioning information from Reb1 to +1 nucleosomes, we turned to classical mononucleosome assays. We generated mononucleosomes with a long linker DNA on one side from a promoter (of gene yGL167c) that was selected based on INO80 and Reb1 occupancy measured by ChIPexo *in vivo*^40^ and clearly improved nucleosome positioning in whole-genome reconstitutions^29^. *In vivo*, the Reb1 site of the yGL167c promoter is 145 bp upstream of the +1 nucleosome dyad (about 72 bp to the 5’ flank of the nucleosome core particle as the distance of this flank to the dyad is about 73 bp) which matches closely the median distance 149 ± 33 bp measured for all Reb1 sites at (median distance to the 5’ flank of 76 ± 33 bp, Fig. 6f). We replaced the +1 nucleosome sequence by a Widom-601 nucleosome positioning sequence^55^ and reconstituted with this construct (Fig. 6c, left) via SGD the *in vivo* promoter nucleosome architecture. Reb1 was added substoichiometrically to reconstituted yGL167c-NCP601 mononucleosomes.

As separation in native polyacrylamide gel electrophoresis could distinguish mono-nucleosomes with and without bound Reb1, we could compare remodeling kinetics with and without Reb1 in the same reaction (Fig. 6c). Kinetics of sliding the initially end-positioned nucleosome to the center were much slower, if at all detectable, in the presence of Reb1 (Fig. 6c,d). As the distance between bound Reb1 and the 601-nucleosome was as *in vivo* and therefore, probably corresponded to the steady state distance set by INO80, we prepared and assayed in the same way a second construct (yGL167c-20-NCP601, Fig. 6c) with additional 20 bp of DNA inserted in the yGL167c promoter. This end-positioned 601-nucleosome was clearly moved towards the Reb1 barrier by INO80 (Fig. 6c), but again at a slower rate compared to sliding this nucleosome to the center in the absence of Reb1 (Fig. 6d).

We asked next, whether decreased sliding kinetics were caused by inhibition or by decoupling of ATPase activity. Notably, most INO80 mutations that abrogate nucleosome sliding, such as the HQ1/2 or Arp5 mutations, still showed robust ATPase activity^43,46^. In contrast, INO80 ATPase assays in the presence of yGL167c-NCP601 mononucleosomes showed about twofold decreased ATPase activity upon addition of Reb1 compared to reactions without Reb1 (Fig. 6e). This was not a general effect of Reb1 in this assay as the HMGII complex as well as the Ino80^ΔN^ INO80 mutant complexes with point mutations in the HMG box of Nhp10 or lacking the N-terminal region of Ino80 and the Nhp10-module, respectively (Fig. 2c, Supplementary Fig. 1h), did not show a reduction of ATPase activity upon Reb1 addition (Fig. 6e), while the ATPase activity of the ΔNhp10 INO80 mutant complex was still regulated by Reb1. The detailed mechanism of this intriguing allosteric communication across a distance of more than 70 bp linker DNA awaits further structural studies. However, based on the regulatory role of the N-terminal region of Ino80 even in the absence of the Nhp10 module, we cautiously speculate that it might serve not only as a binding platform for Nhp10, but that it stimulates the activity of INO80 in absence of Reb1 possibly via restricting the dynamics of the Arp8 module.

Taken together, we concluded that Reb1 binding to its cognate promoter sites regulates INO80 activity allosterically by inhibition through interaction via the N-terminal region of Ino80 that is modulated by the Nhp10 module subunits. The multi-subunit architecture of INO80 relays thereby positioning information between Reb1 and +1 nucleosomes, adjusts the +1 nucleosome to its *in vivo*-like position and programs thereby genic regions for formation of nucleosome arrays (Fig. 6f).

### INO80 integrates information from DNA shape/mechanics and Reb1 at promoters

A Reb1 site at a distance to a nucleosome position corresponds to an input of DNA sequence information, mediated by its bound cognate factor Reb1, compared to the input of DNA shape/mechanics features. Therefore, we asked if and how INO80 serves as an information processing hub and integrates such different information input into resulting nucleosome positions.

First, we asked if promotors with Reb1 sites at all contained DNA shape information leading to +1 nucleosome positioning by INO80 on its own in the absence of Reb1. Maybe promoter regions had evolved such that +1 nucleosome positions were either directly encoded via DNA shape/mechanics or indirectly via GRF sites. We compared nucleosome positioning by INO80 in the absence of Reb1 at Reb1 site-containing promoters with positioning at an equal number of promoters lacking any GRF sites. As INO80 was able to position *in vivo*-like +1 nucleosomes on its own at both types of promoter regions (Fig. 7a), we concluded that both types contained +1 nucleosome position DNA shape/mechanics information in their genomic sequence.

**Figure 7.**
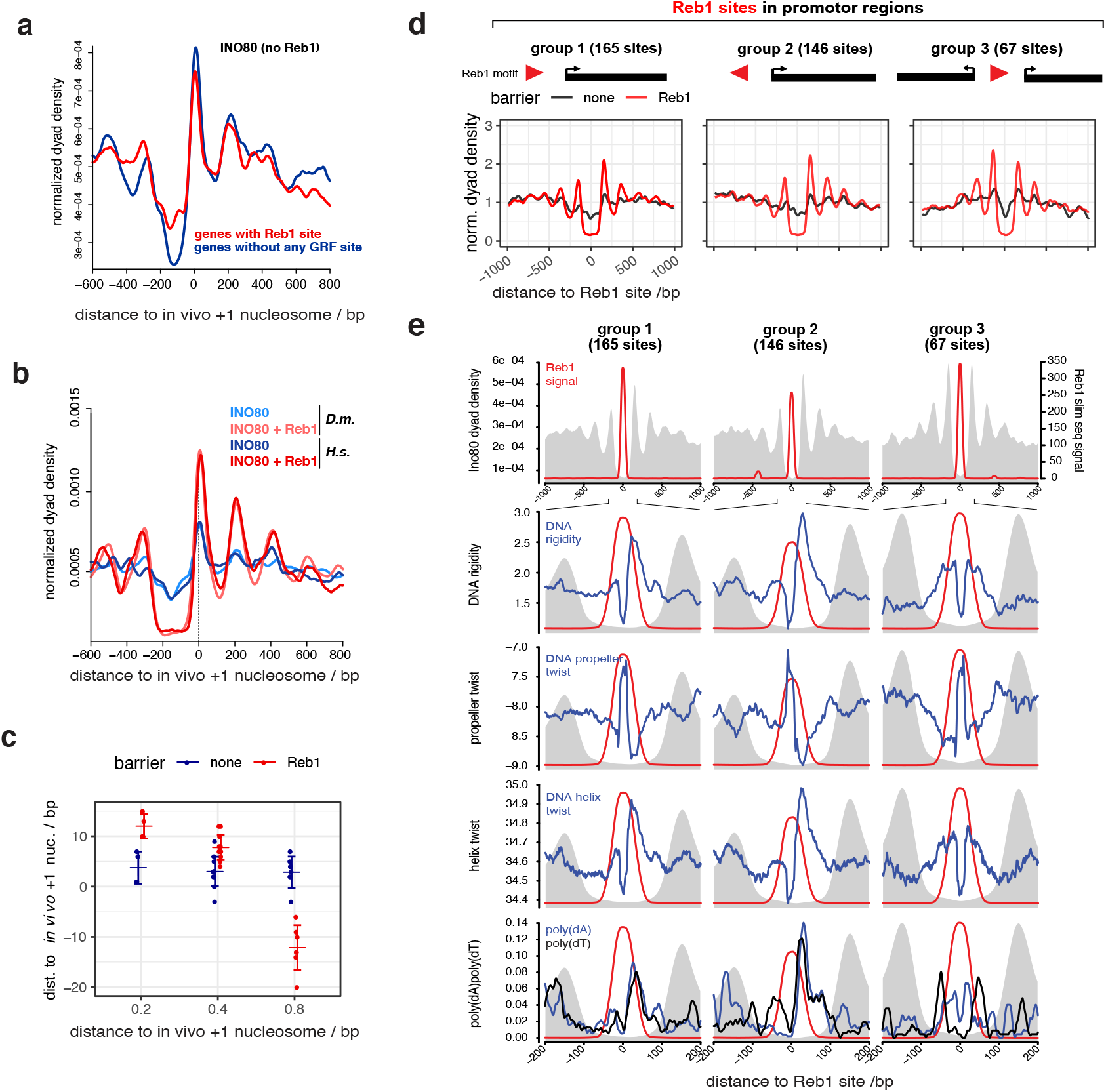
INO80 synergistically integrates nucleosome positioning information from DNA shape and Reb1 barriers. **a** Composite plots as in Figure 6b but for SGD chromatin with recombinant human histones at 0.4 histone-to-DNA mass ratio incubated with recombinant *S. cerevisiae* WT INO80 plotted for either genes with promoter Reb1 sites only (as red shading in Figure 6a) or for a randomly selected but similar number of genes with no GRF sites (Abf1, Rap1, Mcm1, Cbf1 ^70^) in their promoters. **b** As panel a but for merged replicates comparing SGD chromatin with embryonic fly (*D. m.*) or recombinant human (*H. s.*) histones, ± 20 nM Reb1 and only for genes with promoter Reb1 sites. **c** Distributions of distances between +1 nucleosome positions at Reb1-site containing promoters reconstituted by incubation of SGD chromatin with the indicated histone-to-DNA mass ratio with recombinant *S. cerevisiae* WT INO80 in the presence (Reb1) or absence (none) of 20 nM Reb1. **d** As Figure 6b, but aligned at Reb1 sites of the indicated groups and with recombinant *S. cerevisiae* WT INO80 ± 20 nM Reb1. **e** Reb1 site-aligned composite plots for genes groups as in panel d, from top to bottom: positions of Reb1 site PWM motifs, Reb1 site motifs and DNA rigidity, Reb1 sites and propeller twist DNA shape features and Reb1 motifs and positions of poly(dA) or poly(dT) elements (> 6 homopolymeric stretches). Grey background in all panels shows composite plot of MNase-seq data as in panel d.

Second, we asked if the additional information of bound Reb1 at the promoters with Reb1 site was synergistic, antagonistic or neutral to the DNA shape/mechanics-guided positioning by INO80. Comparing nucleosome positioning by INO80 at Reb1 site-containing promoters with versus without Reb1 showed that the Reb1 information mainly synergized with the DNA shape/mechanics information and led to very similar positions but, in keeping with the outcome of the Reb1 titration (Fig. 6a,b), to higher +1 nucleosome peaks and more pronounced NDRs (Fig. 7b). Quantification of the differences in resulting peak positions with vs. without Reb1 showed that +1 nucleosome peaks differed on average only by 6 ± 3 bp for SGD chromatin with histone-to-DNA mass ratios of 0.2 or 0.4, which was within the experimental error of our reconstitutions (Fig. 7c). For higher assembly degrees with a histone-to-DNA mass ratio of 0.8, the difference was 15 ± 5 bp, which was due to nucleosome positioning closer to Reb1 with increasing histone density, while the +1-nucleosome position as determined by INO80 on its own via DNA shape/mechanics was hardly affected by variations in nucleosome density. Nonetheless, high density affected peak heights, which is discussed, together with the effects of density on nucleosome distance to barrier, in the accompanying paper (Oberbeckmann & Niebauer et al.) in the context of our remodeler ruler concept. Here, we concluded that genome sequence evolved a DNA shape/mechanics signal downstream of a Reb1 site in direction of transcription so that nucleosome positioning by INO80 either guided by DNA shape/mechanics or by Reb1 leads to very similar +1 nucleosome positions at low or medium nucleosome density. Note that promoter Reb1 sites are situated *in vivo* within NDRs^56^, which, by definition, represent regions of locally low nucleosome density.

Third, we noted that the synergism between DNA shape/mechanics- and Reb1-guided nucleosome positioning by INO80 only applied to the +1 nucleosome in direction of transcription, but not to the −1 nucleosome, as we observed in our reconstitution experiments in in vivo-like differences between the respective MNase-seq peak heights (Fig. 7b).

To assess this point more clearly and to ask if orientation of the intrinsically asymmetric Reb1 site further affected nucleosome positioning, we grouped Reb1 site-containing promoters according to the Reb1 site orientation relative to neighboring genes (groups 1 to 3, Fig. 7d). Reb1 site-aligned MNase-seq data composite plots averaged over genes within these groups showed that peak heights and array generation were more pronounced in direction of transcription but independent of Reb1 site orientation. This further supported our conclusion that synergistic DNA shape/mechanics information evolved next to Reb1 sites only in places where a +1 nucleosome becomes positioned that plays the well-known role in regulation of transcription initiation^4,28^. Accordingly, promoters in groups 1 to 3 showed distinct asymmetrical DNA shape/mechanics features and strand-specific poly(dA:dT) prevalence in the direction of transcription (Fig. 7e). Thus, these data suggest that INO80-mediated +1 nucleosome positioning is symmetrically guided by Reb1 as orientation of the Reb1 site did not matter (group 1 vs. 2, Fig. 7d). Importantly, however, our analysis revealed that Reb1 sites at promoters evolved synergistically with DNA shape/mechanics features, which explains the observed peak height asymmetry (groups 1 and 2) or symmetry (group 3) of nucleosome patterns depending on the DNA shape/mechanics feature distribution in the genome (Fig. 7e). The deviations in +1 nucleosome positions between DNA shape/mechanics-versus Reb1-guided positioning (Fig. 7c) in response to nucleosome density suggest that Reb1-guided positioning is either dominant or that Reb1-guided positioning is still equally effective at high density while DNA shape/mechanics-guided positioning is impaired. In the accompanying paper (Oberbeckmann & Niebauer et al.) we show that the latter is the case.

### DNA ends are potent barriers for INO80 nucleosome positioning

Having established a synergy between DNA shape/mechanics and Reb1 sites at gene promoter regions, we asked whether we can uncouple barrier-mediated positioning from a promoter sequence context. To test this idea, we analyzed nucleosome positioning at all *in vivo* mapped genomic Reb1 sites (Fig. 8a,b). Consistent with our findings above, we observed symmetrical nucleosome arrays around all Reb1 sites (Fig. 8b, top right) suggesting that barrier-mediated positioning can occur independently of other DNA sequence features. In light of this, we considered that INO80 may align nucleosomes also to different barrier types as long as they represented a clear alignment point. In our search of the minimalistic system that provides nucleosome positioning information, we wondered if simply a DNA end could constitute a barrier. Notably, INO80 has been involved in DNA damage response signaling upon DNA double strand breaks (DSBs) *in vivo*^57^. In principle, such as scenario was already tested in classical mononucleosome sliding assays as these automatically involve two DNA ends.

**Figure 8.**
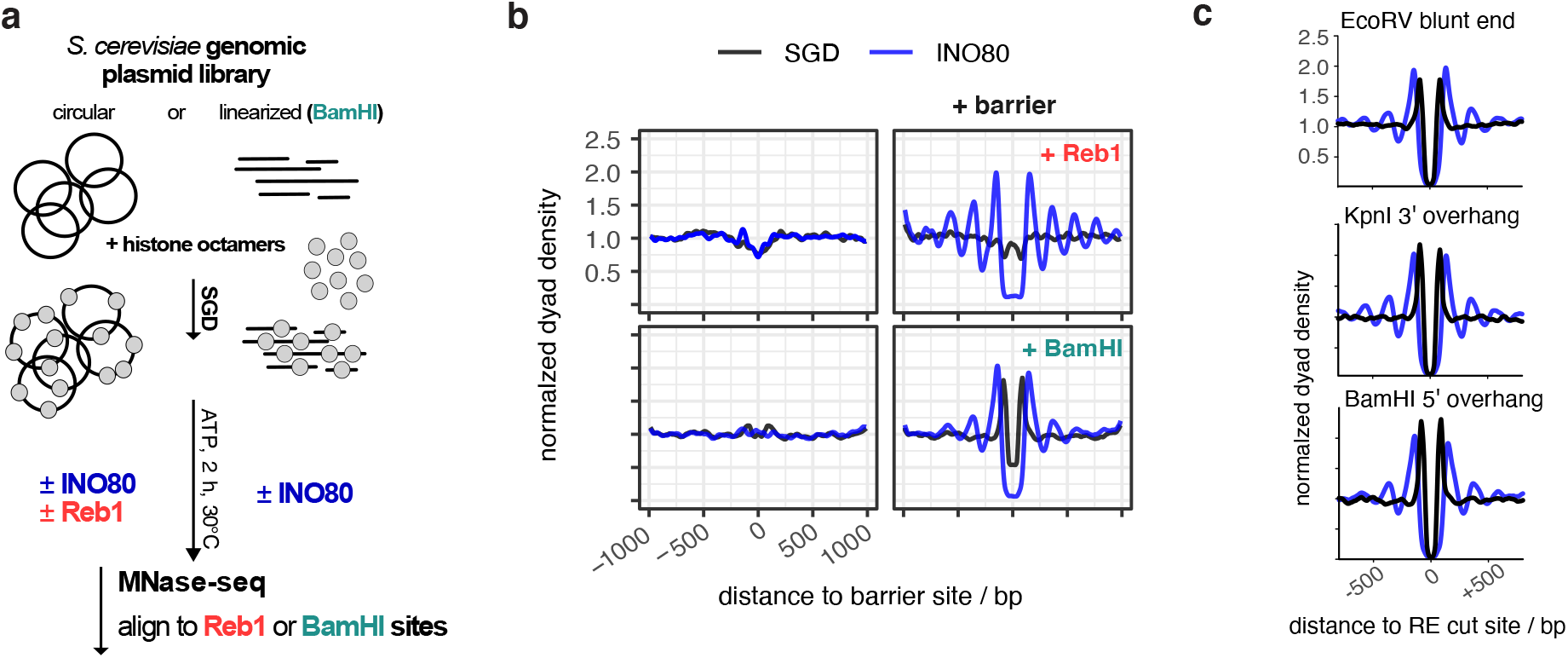
DNA ends are potent barriers for nucleosome positioning by INO80. **a** Overview (analogous to Figure 2a) of reconstitution with circular versus RE-precleaved plasmid libraries. **b** Composite plots of BamHI-site aligned versus anti-Reb1 SLIM-ChIP-defined Reb1 sites aligned MNase-seq data for: top, SGD prepared with circular plasmid library and incubated without (SGD) or with recombinant *S. cerevisiae* WT INO80 (INO80), and bottom: as top but with BamHI-precleaved library if indicated (+ BamHI). **c** As panel b, but for SGD chromatin with plasmid libraries pre-cleaved with the indicated RE and data aligned at the indicated RE cut sites. Strong peaks flanking cut RE sites in SGD chromatin without INO80 remodeling reflected an MNase-seq bias. Due to the pre-cleavage, the probability that a mono-nucleosomal fragment flanking the cut site is released by MNase is increased.

However, effects there may have been due to the comparatively short length of template DNA and to the presence of two DNA ends at the same time. Our genome-wide system allowed us to test the effect of one-sided DNA ends in the context of very long DNA. We introduced double stranded DNA ends at fortuitous locations, i.e., without likely evolutionarily shaped context, throughout the *S. cerevisiae* genome via restriction enzyme (RE) digest of the plasmid library prior to SGD reconstitution (Fig. 8a). As expected, SGD chromatin neither with nor without remodeling by INO80 showed distinct nucleosome patterns at uncleaved BamHI sites (Fig. 8b, bottom left). However, strong and symmetrical arrays were aligned at cut sites by INO80 (Fig. 8b, bottom right). The same was true for other REs that generated different kinds of DNA ends (Fig. 8c). We concluded that all three kinds of DNA ends (blunt, 3’ or 5’ overhang) were strong nucleosome positioning barriers for INO80.

## Discussion

In this study, we identified and probed the fundamental molecular determinants by which ATP-dependent chromatin remodelers position nucleosomes across the genome. An integrated approach combining fully-recombinant, *de novo* whole-genome reconstitutions, high-resolution structural information, and PCA/clustering analysis revealed that the INO80 complex processes DNA sequence information, both via readout of a distinct DNA shape/mechanics signature motif, as well as, via alignment against a DNA sequence specific barrier factor like Reb1 or at DSBs. INO80’s multi-subunit architecture integrates the readout of different positioning information, contributes through its mechanism its own information and determines thereby how this is translated into positions of +1 and other nucleosomes (Fig. 9).

**Figure 9.**
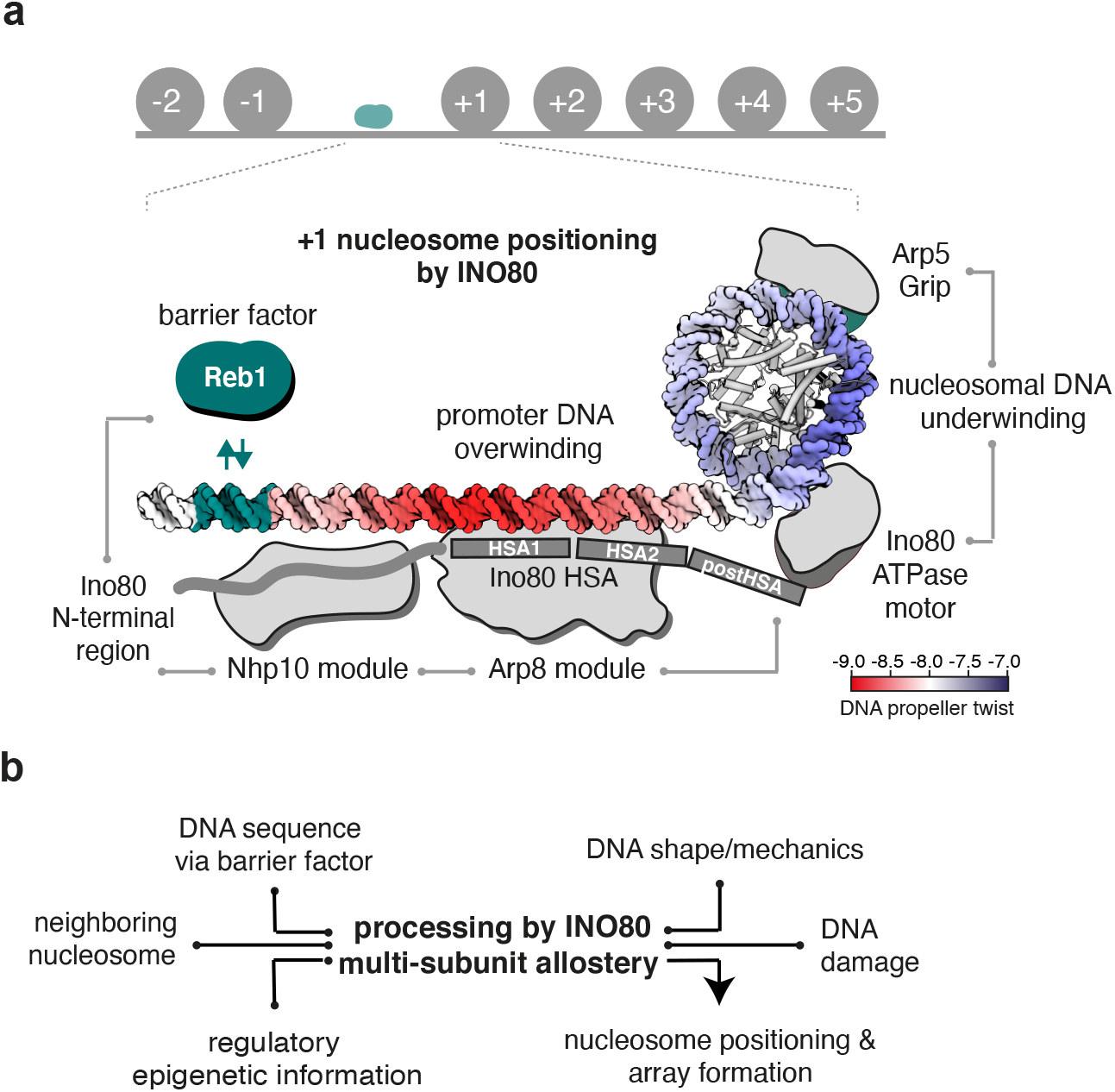
Model of +1 nucleosome positioning by INO80. **a** INO80 synergistically processes genomic information derived from DNA shape/mechanics as well as DNA sequence motifs bound by GRF Reb1 to position +1 nucleosomes. Structural data^43,46^, biochemical^47^ and ChIP-exo mapping^40^ suggest a binding architecture of INO80 at +1 nucleosomes that is fully consistent with the identified positioning information and mechanism. Promoter DNA overwinding and nucleosomal DNA underwinding is derived from the direction of DNA translocation by the Snf2-type ATPase of INO80^43^. Allosteric communication is indicated by grey lines. **b** Signal integration and processing by multi-subunit allostery within INO80 leads to nucleosome positioning and array formation. Epigenetic information such as histone marks are expected to provide an additional layer of regulatory input, e.g., in response to the physiological state of the cell.

Although the pivotal role of remodelers in chromatin organization and their dependency on DNA sequences has been recognized^29,31,58^, nucleosome positioning sequences (NPSs) were usually defined as sequences of “intrinsic” positioning by SGD driven solely by histone octamer-DNA interactions, as illustrated by the Widom-601 NPS^55^. PCA/clustering analysis enabled us now to reassess these classical SGD-NPSs and to identify a new kind of NPS. We find that SGD-NPSs correspond to distinct DNA-sequence dependent shape/mechanics profiles, while nucleosome positioning by a remodeler like INO80 corresponds to a different shape/mechanics profile. Therefore, we identified the latter as INO80-NPSs.

Respective remodeler-NPSs are likely to exist for other remodelers and it will be interesting where they evolved in genomes. The mere observation that INO80 and RSC remodelers generate different nucleosome positions, despite working on the same histone octamers and DNA sequences, suggested previously^29, 59^ that remodelers do not just allow histone octamers to occupy their thermodynamically preferred positions (otherwise different remodelers would generate the same positions), but that remodelers, as demonstrated in this study, read genomic information, actively override octamer preferences and shape the positioning landscape in a remodeler-specific way. In analogy to the “genomic code for nucleosome positioning”, i.e. the proposed evolution of SGD-NPSs, evolved remodeler-NPSs would implement a “remodeler code for nucleosome positioning” as proposed earlier^59^. We abstain from adding another “code” to the troubled epigenetics discussions but point out the conceptual analogy.

Importantly, we go here beyond a mere correlation between INO80-NPSs and DNA shape/mechanics profiles. The causal mechanistic link was directly established by tuning the INO80 DNA shape/mechanics readout via targeted INO80 mutations. Informed by high-resolution structures, we found independently that on the one hand mutation of Ino80-HSA-DNA contacts more than −100 bp away from the nucleosome dyad caused altered nucleosome positioning patterns, while on the other hand unbiased PCA/clustering analysis revealed also altered DNA shape/mechanics features right in the same region. Together, our results provide strong evidence for a readout of these DNA shape/mechanics features. Moreover, we observed altered processing of DNA shape/mechanics features at the −55 bp region between the Ino80 core ATPase motor and the Arp5 grip, suggesting a critical role of DNA shape/mechanics in regulating the build-up of DNA strain during the core mechanism of nucleosome translocation^43,47,48^. Intriguingly, the effects at both regions are coupled via two allosteric communication pathways of possibly equal importance: on the protein side, linker DNA recognition by the Arp8 module is coupled to the activity of the Ino80 ATPase motor of the core module via the extended helical configuration of the HSA and postHSA domains^46^. On the DNA side, DNA shape/mechanics features at the histone-bound −55 bp region are most likely coupled to DNA shape/mechanics features at the DNA linker −100 bp region in the context of over- and underwinding of DNA in front and behind the Ino80 ATPase motor^38,43^. More generally, our data illustrates a regulatory circuitry comprising a two-way relationship between a protein factor working on DNA and DNA properties feeding back to the protein factor. Overall, INO80-NPSs represent the nucleosome positioning information that emerges from the combination of DNA, histones, and the active interpretation via the allosteric communication within the remodeler.

For these reasons, the DNA shape/mechanics readout by INO80 importantly expands the scope of recently discussed DNA shape contributions. DNA shape was mostly studied in the context of “static” DNA binding, e.g., by transription factors and GRFs^60–62^. In contrast, INO80 dynamically reads and interprets DNA shape/mechanics while tracking along DNA in an ATP-dependent manner. Thereby, INO80 actively probes the mechanical properties of DNA. Thus, this read out of genome information is expected to serve as a role model for other factors that translocate along DNA or also RNA, like other remodelers, helicases, cohesins or polymerases. For example, RNA polymerase I was suggested to read the DNA bend at its promoters^63^ and RNA polymerase II may recognize its promoters via structural DNA features (bending, meltability, flexibility) rather than via classical consensus sequences^64^. As these structural properties are redundantly linked to DNA sequence, we propose that readout of such DNA structural properties may be common if factors deal with a wide range of genomic regions.

As alternative DNA sequence signals, there is DNA sequence information of classical consensus motifs for specific binding by cognate factors. GRFs are well-known to program +1 nucleosome positioning and formation of genic nucleosome arrays *in vivo*^26,34,65^. In light of our finding that DNA ends are also potent nucleosome positioning barriers, it is tempting to speculate that remodelers involved in DNA damage response, such as INO80^57^, may generate regular nucleosome arrays as a licensing platform at DSBs *in vivo*.

The mechanism by which remodelers generate arrays at barriers, i.e., read positioning information via an alignment mechanism, remained largely unknown. This study reveals that nucleosome positioning by INO80 is actively regulated by Reb1 at promoter sites through an interaction with the N-terminal region of Ino80 (Fig. 9a). Intriguingly, Reb1 decreased not only nucleosome sliding, but also inhibited ATPase activity of INO80, even at a distance of −145 bp between the cognate Reb1 site and the dyad of the +1 nucleosome. In contrast, DNA linker length sensing by INO80 at DNA ends uncouples a decrease in mononucleosome sliding from its robust stimulation of ATPase activity^46,48^. Consequently, GRFs might represent a different kind of regulatory barrier compared to DSBs, at least in the absence of the DNA repair machinery. In the accompanying study (Oberbeckmann & Krietenstein et al.), we identify the Arp8-module and the Nhp10 module as a multi-layered ruler element which measures and sets nucleosome arrays differently in respect to Reb1 sites, DNA ends and neighboring nucleosome. Taken together, our findings lead to a model how regulation of nucleosome sliding direction bias upon interaction with a barrier can lead to stable nucleosome positioning and array formation. The multi-subunit architecture of INO80 functions similarly to a relay: INO80 receives input via its Arp8 and Nhp10 modules and communicates this information allosterically towards the ATPase of the INO80 core, where it is translated into a nucleosome position (Fig. 9b).

The exact +1 nucleosome position impacts transcription regulation, e.g., it differs between repressed and activated promoters and influences TSS selection^4,11,28,66^. In this study, we show that these positions are robustly encoded in the genome in two ways, i.e., both by DNA shape/mechanics features and corresponding distances to the Reb1 site. Nucleosome positioning next to Reb1 did not require DNA shape/mechanics features as it also worked symmetrically on the other side even if there was no evolved promoter. Importantly, however, in context of promoter regions, we identify a co-evolved synergy between DNA shape/mechanics signatures and Reb1 binding sites, leading to asymmetric +1 nucleosome positioning, as measured by MNase-seq peak heights. This synergy provides not only robustness, but also an inroad to regulation. For example, we show that Reb1-mediated positioning is altered in response to nucleosome densities. Thus, we propose that regulation of nucleosome density at promoters, e.g., via the local activity of RSC, the major nucleosome-evicting remodeler in yeast^23^, may result in regulation of +1 nucleosome positions. With high RSC activity, local promoter nucleosome density is low and +1 nucleosome positioning by INO80 coincides for DNA shape/mechanics- and Reb1-information input. Upon low RSC activity, nucleosome density is high and INO80 disregards the shape/mechanics signal and places the +1 nucleosome closer to Reb1, which corresponds to the more upstream +1 nucleosome position implicated in repressed promoter states.

By genome wide biochemistry, this study reveals that a minimal set of information, comprising genomic DNA sequences, globular histones, and the molecular machinery of the remodeler, is sufficient to explain the placement and regulation of nucleosomes at their *in vivo* +1 positions for many promoters where appropriate DNA shape/mechanics signatures evolved. The identified mechanism of active information processing (Fig. 9b) provides allosteric control and versatile means for selective regulation, e.g., by epigenetic information such as histone modifications and variants as well as by the presence of sequence-specific factors such as transcription factors and pioneer factors. Signal integration of genome information from DNA shape/mechanics and sequence specified GRF binding by the multi-subunit architecture of INO80 exemplifies such principles. In the accompanying paper (Oberbeckmann & Niebauer et al.), we show how information from GRFs, DNA ends and positioned nucleosomes can be propagated into regular nucleosome arrays and how this process is regulated by remodeler rulers and nucleosome density. Collectively, this makes ATP dependent remodelers the fundamental information processing hub for nucleosome positioning and thereby the primary architects of the first level of chromatin organization.

## Acknowledgments

We thank Stefan Krebs and Helmut Blum at the Laboratory for Functional Genome Analysis (LAFUGA, Gene Center, LMU München) for high throughput sequencing, and Sigurd Braun for access to and help with the plating robot. We thank Marianne Schwarz and Jens Michaelis for helpful discussions and earlier experiments using recombinant histones and endogenously purified INO80 not reported in this paper. This study was funded by the German Research Foundation (SFB1064 to P.K. and K.P.H. and Gottfried Wilhelm Leibniz-Prize to K.P.H.), the European Research Council (ERC Advanced Grant “INO3D” to K.P.H.), and the NIH (grant R35GM130376 to R.R.) and HFSP (grant RGP0021/2018 to R.R.). N.K. is supported by HFSP grant LT000631/2017-L.

## Author contributions

Conceptualization: EO, NK, TS, RR, KPH, PK, SE; Data curation: EO, NK, VN; Formal analysis: EO, NK, VN, YW, TS; Funding acquisition, Project administration, Supervision: KPH, PK, SE; Investigation: EO, NK, VN, KS, MM, YW, TS, SE; Methodology: EO, NK, VN, KS, RR, TS, PK, SE; Validation: EO, NK, VN, KS, YW, RR, TS, PK, SE; Visualization: EO, NK, VN, YW, TS, PK, SE; Writing original draft: EO, NK, PK, SE; Writing – review & editing: EO, NK, VN, TS, RR, KPH, PK, SE.

## Methods

### Organisms Embryonic *D. melanogaster* histones, whole-genome plasmid libraries and salt gradient dialysis

#### Embryonic *D. melanogaster* histone purification

The preparation of embryonic *D. melanogaster* histones octamers was carried out as described before^1,2^. In brief, 50 g of 0-12 hours old *D. melanogaster* embryos (strain OregonR) were dechorionated in 3 % sodium hypochlorite, washed with dH20 and resuspended in 40 mL lysis-buffer (15 mM K·HEPES pH 7.5, 10 mM KCl, 5 mM MgCl_2_, 0.1 mM EDTA, 0.5 mM EGTA, 1 mM DTT, 0.2 mM PMSF, 10 % glycerol). Embryos were homogenized (Yamamoto homogenizer), filtered through cloth and centrifuged at 6,500 *g* for 15 min. Nuclei (brownish light pellet) were washed 3 times with 50 mL sucrose-buffer (15 mM K·HEPES pH 7.5, 10 mM KCl, 5 mM MgCl_2_, 0.05 mM EDTA, 0.25 mM EGTA, 1 mM DTT, 0.2 mM PMSF, 1.2 % sucrose) and resuspended in 30 mL sucrose-buffer containing 3 mM CaCl_2_. To obtain mononucleosomes, nuclei were incubated for 10 min at 26 °C with 6250 Units MNase (Sigma-Aldrich). Reaction was stopped with 10 mM EDTA, nuclei were pelleted and resuspended in 6 mL TE (10 mM Tris·HCl pH 7.6, 1 mM EDTA) containing 1 mM DTT and 0.2 mM PMSF followed by 30 to 45 min of rotation at 4 °C. Nuclei were octamers were eluted with 2 M KCl, concentrated and stored in 50 % glycerol and 1x Complete (Roche) protease inhibitors without EDTA at −20 °C.

#### Whole-genome plasmid library expansion

The *S. cerevisiae* genomic plasmid library (pGP546) was originally described by Jones et al.3 and purchased as a clonal glycerol stock collection from Open Biosystems. Library expansion was carried out via a Singer ROTOR plating machine (Singer Instruments) (8-12 rounds, 3 replicas). After 16 hours, colonies were combined into 3×2 L of LB medium containing 50 μg/mL kanamycin and grown for 4 hours. Cells were harvested and subjected to Plasmid Giga Preparation (PC 10 000 Kit, Macherey&Nagel).

#### Salt gradient dialysis (SGD)

For low, medium and high assembly degrees, 10 μg of plasmid library DNA (*S. cerevisiae*, *S. pombe* or *E. coli*) was mixed with ~2, 4 or 8 μg of *Drosophila* embryo histone octamers, respectively, in 100 μl assembly buffer (10 mM Tris·HCl, pH 7.6, 2 M NaCl, 1 mM EDTA, 0.05 % IGEPAL CA630, 0.2 μg BSA). For reconstitutions with precleaved DNA (Fig. 8), the plasmid library was digested with the respective restriction enzyme and purified by phenol extraction/ethanol precipitation prior to SGD. Samples were transferred to Slide-A-lyzer mini dialysis devices, which were placed in a 3 L beaker containing 300 mL of high salt buffer (10 mM Tris·HCl pH 7.6, 2 M NaCl, 1 mM EDTA, 0.05 % IGEPAL CA630, 14.3 mM β-mercaptoethanol), and dialyzed against a total of 3 L low salt buffer (10 mM Tris·HCl pH 7.6, 50 mM NaCl, 1 mM EDTA, 0.05 % IGEPAL CA630, 1.4 mM β-mercaptoethanol) added continuously via a peristaltic pump over a time course of 16 h while stirring. β-mercaptoethanol was added freshly to all buffers. After complete transfer of low salt buffer, samples were dialyzed against 1 L low salt buffer for 1 h at room temperature. DNA concentration of the SGD chromatin preparations was estimated with a DS-11+ spektrophotometer (Denovix) and could be stored at 4 °C for several weeks. To estimate the extent of the assembly degree, an aliquot of the sample was subjected to MNase digestion (as described below) for MNase-ladder read out.

#### Expression and purification of INO80 complex and respective mutants

Coding sequences for *S. cerevisiae* Ino80 (2xFlag), Rvb1, Rvb2, Arp5-His, Ies6 (pFBDM_1) and Actin, Arp4, Arp8, Taf14, Ies2, Ies4, Ies1, Ies3, Ies5 and Nhp10 (pFBDM_2) were subcloned into pFBDM vectors^4^ and sequence verified by Sanger Sequencing (GATC Services at Eurofins Genomics). Bacmids of both vectors were generated using DH10 multibac cells^5^. Baculoviruses were generated in *Spodoptera frugiperda* (SF21) insect cells (IPLB-Sf21AE). *Trichoplusia ni* High Five (Hi5) insect cells (BTI-TN-5B1-4 Invitrogen) were co-infected with two baculoviruses 1/100 each. After 60 h cultivation at 27 °C, cells were harvested by centrifugation. For purification of the INO80 complex, cells were resuspended in lysis buffer (50 mM Tris·HCl pH 7.9, 500 mM NaCl, 10 % glycerol, 1 mM DTT, SIGMAFAST™ protease inhibitor cocktail), sonified (Branson Sonifier, 3x 20 s with 40 % duty cycle and output control 3-4) and cleared by centrifugation (Sorvall Evolution RC, SS34 rotor, 15,000 *g*). The supernatant was incubated for 1 h with anti-Flag M2 Affinity Gel (Sigma-Aldrich) and centrifuged for 15 min at 1,000 *g* and 4 °C. The anti-Flag resin was washed with buffer A (25 mM K·HEPES pH 8.0, 500 mM KCl, 10 % glycerol, 0.025 mM IGEPAL CA630, 4 mM MgCl_2_, 1 mM DTT) and buffer B (25 mM K·HEPES pH 8.0, 200 mM KCl, 10 % glycerol, 0.02 mM IGEPAL CA630, 4 mM MgCl_2_, 1 mM DTT).

Recombinant INO80 complex was eluted with buffer B containing 1.6 mg Flag Peptide (Sigma-Aldrich). Anion exchange chromatography (MonoQ 5/50 GL, GE Healthcare, Buffer: 25 mM K·HEPES pH 8.0, 4 mM MgCl_2_, 1 mM DTT) using a linear KCl gradient 200mM-1000mM) and, if required, size exclusion chromatography (Superose 6, 10/300 GL, 25 mM K·HEPES pH 8.0, 200mM, 4 mM MgCl_2_, 1 mM DTT) was used for further purification which resulted in a monodisperse INO80 complex (Figure S1A,B,E). Using standard cloning techniques, three INO80 (2xFlag) HSA domain mutants (HQ1, HQ2, HQ1/2; Figure 2C, S1E), one N-terminal deletion mutant (Ino80^ΔN^, deletion of the first 461 amino acids of the N-terminus of Ino80) and two INO80 (2xFlag) Nhp10 module mutants (ΔNhp10 (INO80 complex without Ies1, Ies3, Ies5 and Nhp10 but with Ino80 N-terminus) and HMGII (Figure 2C, S1E) pFBDM vectors were generated and integrated into baculoviruses using MultiBac Technology as described above. Expression and purification of mutant INO80 complexes was essentially carried out as WT INO80 complex purification. The INO80 core complex from *Chaetomium thermophilum* (equivalent to the *S. cerevisiae* N-terminal deletion mutant) was essentially purified as described in ^6^.

#### Genome-wide remodeling reaction

All remodeling reactions were performed at 30 °C in 100 μL with final buffer conditions of 26.6 mM Na·HEPES pH 7.5, 1 mM Tris·HCl pH 7.6, 85.5 mM NaCl, 8 mM KCl, 10 mM ammonium sulfate, 10 mM creatine phosphate (Sigma-Aldrich), 3 mM MgCl_2_, 2.5 mM ATP, 0.1 mM EDTA, 0.6 mM EGTA, 1 mM DTT, 14 % glycerol, 20 ng/μl creatine kinase (Roche Applied Science). Remodeling reactions were started by adding 10 μL SGD chromatin corresponding to ~ 1 μg DNA assembled into nucleosomes and terminated by adding 0.8 Units apyrase (NEB) followed by incubation at 30 °C for 30 min. Independent replicates of remodeling reactions refer to independent SGD chromatin preparations. The experimental conditions for each sample are detailed in Supplementary Data 1.

#### MNase-seq

After apyrase addition, remodeling reactions were supplemented with CaCl2 to a final concentration of 1.5 mM and digested with 100 Units MNase (Sigma) to generate mostly monoucleosomal DNA. 10 mM EDTA and 0.5 % SDS (final concentrations) were added to stop the MNase digest. After proteinase K treatment for 30 min at 37 °C, samples were ethanol precipitated and electrophoresed for 1.5 - 2 h at 100 V using a 1.5 % agarose gel in 1x Tris-acetate-EDTA (TAE) buffer. Mononucleosome bands were excised and purified with PureLink Quick Gel Extraction Kit (ThermoFisher Scientific). For library preparation, 10-50 ng of mononucleosomal DNA was incubated with 1.25 Units Taq polymerase (NEB), 3.75 Units T4 DNA polymerase (NEB) and 12.5 Units T4-PNK (NEB) in 1x ligation buffer (B0202S, NEB) for 15 min at 12 °C, 15 min at 37 °C and 20 min at 72 °C. To ligate NEBNext Adaptors (0.75 μM final concentration, NEBNext Multiplex Oligos Kit) to the DNA, samples were incubated with T4 DNA ligase (NEB) at 25 °C for 15 min, followed by incubation with 2 Units USER enzyme (NEB) for 10 min at 37 °C. Fragments were purified using 2 volumes AMPure XP beads (Beckman Coulter) and amplified for 8-10 cycles using NEBNext Multiplex Oligos, Phusion High-Fidelity DNA Polymerase (1 U, NEB), deoxynucleotide solution mix (dNTP, 2.5 mM, NEB) and Phusion HF Buffer (1x, NEB). The following protocol was applied for amplification: 98 °C for 30 s, 98 °C for 10 s, 65 °C for 30 s, 72 °C for 30 s with a final amplification step at 72 °C for 5 min. DNA content was assessed by using Qubit dsDNA HS Assay Kit (Invitrogen). PCR reactions were applied to an 1.5 % agarose gel, needed fragment length (~270 bp) was excised and purified via PureLink Quick Gel Extraction Kit (ThermoFisher Scientific). DNA was measured again with Qubit dsDNA HS Assay Kit and diluted to a final concentration of 10 nM (calculation based on the assumption that the DNA fragment length is 272 bp, i.e., 147 bp nucleosomal DNA and 122 bp sequencing adaptor). Diluted samples were pooled according to sequencing reads (~6 Mio reads/ sample). The final pool was quantified with BioAnalyzer (Agilent) and analyzed on an Illumina HiSeq 1500 in 50 bp single-end mode (Laboratory for Functional Genome Analysis, LAFUGA, LMU Munich).

#### Expression and purification of human tailless histone octamers

The genes for expression of tailless human histones H2A, H2B and H4 were cloned in pET21b vectors (Merck, Darmstadt, Germany) by blunt-end ligation of genes coding for full-length human histones. The gene coding for human tailless H3 was cloned in a pETM-11 vector (kindly provided by EMBL, Heidelberg, Germany) carrying a N-terminal SUMO-tag by Gibson assembly^7^. The SUMO-tag was removed during octamer assembly. Constructs of tailless histones were designed according to globular domains identified by tryptic digest of full-length histone^8–10^ and comprised the following amino acids: H2A: 13 – 118; H2B: 24 – 125; H3: 27 – 135; H4: 20 – 102. Histones were purified by a combination of inclusion body purification and ion-exchange chromatography, essentially as described previously^11,12^. In brief, histones were expressed in *E. coli* BL21 (DE3) cells (Merck, Darmstadt, Germany) for 2 h after induction with 1 mM IPTG at 37 °C and disrupted under non-denaturing conditions to separate inclusion bodies from lysate. Inclusion bodies were first washed with 1% Triton-X100. Subsequently, inclusion bodies were resuspended in 7 M guanidinium chloride and dialyzed against 8 M urea. Individual histones were purified by cation-exchange chromatography, refolded under low-salt conditions and polished by anion-exchange chromatography. For long-time storage, histones were lyophilized overnight. For octamer reconstitution, histones were resuspended in 25 mM Tris, pH 7.5, 7 M guanidinium chloride, 0.25 mM DTT, mixed at 1.2-fold excess of H2A and H2B and dialyzed against 25 mM Tris·HCl pH 7.5, 2 M NaCl, 0.25 mM DTT overnight. 1 mg/mL SENP2 protease was added after 3 h. The octamer of tailless histones was purified by size-exclusion chromatography using a Superdex 200 16/60 column (GE Healthcare), which separated the octamer from aggregate, H2A/H2B dimers, the SENP2 protease and the SUMO-tag. The purification was analyzed on a 18 % polyacrylamide SDS gel stained with Coomassie (data not shown). The octamer was concentrated to 3.0 mg/mL and stored at −20°C in 50% glycerol.

#### Expression and purification of *S. cerevisiae* Reb1

For genome-wide remodeling reaction *S. cerevisiae* Reb1 was purified exactly as described in^13^. For ATPase and mononucleosome sliding assays Reb1 was purified as follows: Reb1 was amplified from BY4741 genomic *S. cerevisiae* DNA by PCR and cloned into pET21b (Novagen) via InFusion cloning (Clontech) with a Streptavidin tag at the C terminus. Correct sequences were verified via Sanger sequencing (GATC Services at Eurofins Genomics). Expression plasmids were transformed into BL21 (DE3) cd^+^ cells. Three liters of LB medium supplemented with 600 mg/L ampicillin were inoculated with 200 mL pre-culture. Cells were grown at 37 °C to an OD_600_ of 0.6 (WPA CO8000 cell density meter). Induction was carried out by addition of IPTG to a final concentration of 1 mM. Cells were grown overnight at 18 °C, harvested by centrifugation (3,500 rpm, Sorvall Evolution RC) and stored at −80 °C. Cells were resuspended in lysis buffer (50 mM Tris·HCl pH 7.9, 500 mM NaCl, 7 % glycerol, 1 mM DTT, 7 % sucrose and protease inhibitor 1:100), sonicated (Branson Sonifier 250, 5 min at 40-50 % duty cycle and output control 4) and cleared by centrifugation (Sorvall Evolution RC, SS34 rotor, 15,000 *g*). The supernatant was dialyzed over night against 2 L low salt buffer (25 mM K·HEPES pH 8.0, 50 mM KCl, 7 % glycerol, 4 mM MgCl_2_, 1 mM DTT). Heparin chromatography (5 mL column, elution buffer: 25 mM K·HEPES pH 8.0, 1 M KCl, 7 % glycerol, 4 mM MgCl_2_, 1 mM DTT) followed by size exclusion chromatography (Superdex 200 10/300, buffer: 25 mM K·HEPES pH 8.0, 200 mM KCl, 7 % glycerol, 4 mM MgCl_2_, 1 mM DTT) were used for purification. Peak fractions were analyzed by Coomassie SDS-PAGE. Fractions containing Reb1 were pooled, concentrated and stored at −80 °C.

#### Preparation of mononucleosomes with recombinant human octamers

Canonical human histones were provided by The Histone Source – Protein Expression and Purification (PEP) Facility at Colorado State University. Lyophilized individual human histones were resuspended in 7 M guanidinium chloride, mixed at a 1.2-fold molar excess of H2A/H2B and dialyzed against 2 M NaCl for 16 h. Histone octamers were purified by size exclusion chromatography (HILoad 16/600 Superdex 200 column, GE Healthcare) and stored at −20 °C in 50 % glycerol.

We used fluorescein-labeled Widom 601 DNA^14^ with 80 bp extranucleosomal DNA (0N80 orientation) harboring an in vivo ChIP-Exo verified Reb1 binding site^15^ of *S. cerevisiae* gene yGL167c (Reb1 binding motif: TTACCC) 64 or 84 bp distant to the 601 sequence. The DNA template (yGL267c_601) was amplified via PCR, purified by anion exchange chromatography (HiTrap DEAE FF, GE Healthcare) and vacuum concentrated. DNA and assembled histone octamer were mixed in 1.1-fold molar excess of DNA at 2 M NaCl. Over a time-period of 17 h at 4 °C the NaCl concentration was reduced to a final concentration of 50 mM NaCl. Again, anion exchange chromatography was used to purify reconstituted nucleosome core particle (NCP) which were then dialyzed to 50 mM NaCl. NCPs were concentrated to 1 mg/mL and stored at 4 °C.

#### ATPase Assay

As described previously^16^, we applied an NADH-based ATPase assay ^17^ to determine INO80’s ATPase rate. 15 nM INO80 were incubated at 30 °C in a final volume of 50 μl assay buffer (25 mM K·HEPES pH 8.0, 50 mM KCl, 5 mM MgCl_2_, 0.1 mg/mL BSA) with 0.5 mM phosphoenolpyruvate, 2 mM ATP, 0.2 mM NADH and 25 units/mL lactate dehydrogenase/pyruvate kinase (Sigma-Aldrich) to monitor the NADH dependent fluorescence signal in non-binding, black, 384-well plates (Greiner) at an excitation wavelength of 340 nm and an emission wavelength of 460 nm over a 40-min period. We used the Tecan Infinite M1000 (Tecan) plate reader for read out. For all samples, ATPase activity was determined at maximum INO80 WT ATPase activity. ATPase activity was stimulated with 25 nM GL167c-0N80 mononucleosomes with or without equimolar ratios WT Reb1. Using maximal initial linear rates corrected for the buffer blank, we calculated final ATP turnover rates.

#### Mononucleosome sliding assay

Nucleosome sliding activity of INO80 wild type and mutant complexes were monitored on Reb1 site-0N80 mononucleosomes in absence and presence of Reb1. INO80 at a concentration of 10 nM was incubated with 90 nM of Reb1 site-0N80 mononucleosomes in sliding buffer at 26 °C (sliding buffer: 25 mM Na·HEPES pH 8.0, 60 mM KCl, 7 % glycerol, 0.10 mg/mL BSA, 0.25 mM dithiothreitol and 2 mM MgCl2). ATP and MgCl2 at final concentrations of 1 mM and 2 mM, respectively, were added to start the sliding reaction. After 30 s, 60 s, 120 s, 300 s, 600 s, 1800 s and 3600 s the reaction was stopped by adding lambda DNA (NEB) to a final concentration of 0.2 mg/mL. To separate distinct nucleosome species, we applied NativePAGE (NativePAGE Novex 4-16 % Bis-Tris Protein Gels, Invitrogen). The fluorescein-labeled mononucleosomal DNA was visualized by a TyphoonTM FLA 9000 imager.

#### Data Processing

Sequencing data was mapped to the SacCer3 (R64) genome using *bowtie*18. Multiple matches were omitted. After mapping, data was imported into R Studio using GenomicAlignments19. Every read was shifted by 73 bp to cover the nucleosome dyad and extended to 50 bp. Genome coverage was calculated, and aligned to either *in vivo* +1 nucleosome positions20, BamHI cut sites, Reb1 SLIM-ChIP hits21 or Reb1 PWM hits22. Signal was normalized per gene in a 2001 bp window centered on the alignment point.

Heatmaps were sorted either by NFR length (distance between in vivo +1 and −1 nucleosome annotated by calling nucleosomes of *in vivo* MNase-seq data, see below) or by Reb1 binding score. For the latter, Reb1 SLIM-ChIP data (GSM2916407) was aligned to in vivo +1 nucleosome positions and sorted by signal strength in a 120 bp-window 160 bp upstream of every +1 nucleosome. For promotor grouping according to Reb1 site orientation, Reb1 SLIM-ChIP hits which contain a PWM site (± 50 bp) and which are located within 400 bp upstream of in vivo +1 nucleosomes were used. Cluster 1 contains promotors where the Reb1 PWM motif is located on the sense strand and cluster 2, where the Reb1 PWM motif is located on the antisense strand. Cluster 3 contains Reb1 sites at bidirectional promotors.

#### DNA shape and poly(dA:dT) analysis surrounding Reb1 binding sites

The DNA sequence of the yeast genome (SacCer3) was downloaded from *Saccharomyces* Genome Database (SGD) and the DNA shape feature scores (helix twist, propeller twist, minor groove width and electrostatic potential) were calculated for the entire genome using the R package *DNAshapeR* (v1.10.0). Similar to^13^, the resulting DNA shape vectors were smoothed with a 5-bp rollmean. For composite analysis, DNA shape feature specific values were extracted in a window of −2000 to 2000 bp around Reb1 binding sites, oriented with respect to Reb1 motif directionality, and averaged by base pair. Plotted distance around Reb1 features are indicated in respective figures. For the poly(dA:dT) analysis, stretches of 6 nucleotide long polyA (5’-AAAAAA-3’) or polyT (5’-TTTTTT-3’) were identified in the yeast genome using R package *Biostrings* (v2.52.0) and counted. For composite analysis, ploy(dA) or poly(dT) counts were extracted in a window of −2000 to 2000 bp around Reb1 binding sites, oriented with respect to Reb1 motif directionality, and averaged by base pair. Plotted distance around Reb1 features are indicated in respective figures.

#### Identification of TSS +1 nucleosomes

+1 nucleosome positions were called according to ^23^. In more detail, mononucleosomal fragments generated from BY4741 MNase digested chromatin were sequenced on an Illumina Genome analyzer, mapped to the SacCer3 genome with *bowtie* ^18^ and shifted by 73 bp with respect to sequencing read directionality to obtain theoretical nucleosome dyads. The obtained dyad-density counts were smoothed with sliding Gaussian filter (width = 100, mean = 0, SD = 25) and resulting values were sorted by decreasing values. Iteratively, the position with the highest value was added to the list of “dyad centers” and all values for positions within +/-120 bp surrounding the position with the highest value were removed from further analysis. The top 90% of nucleosome dyad centers, by value, constituted the final list of nucleosome positions. Plus 1 nucleosome dyad positions were defined as the nearest nucleosome dyad position to TSS within a window 0 to +500 bp from the TSS, with respect to direction of transcription.

#### Genome-wide principal component and DNA shape analysis of nucleosomes

For PCA and DNA shape analysis, mononucleosomes were sequenced in 50 bp paired-end mode on an Illumina HiSeq1500. If not stated otherwise, functions were called with default parameters. Read pairs were aligned using bowtie2 (version 2.2.9) with options “-X 250 --no-discordant --no-mixed --no-unal”. Only unique matches were kept, and orphaned mates removed. Nucleosomes were called on each sample using bioconductor/nucleR (2.16.0) on nucleosomal fragments defined by paired reads as follows: fragments were processed with trimming to 40 bp around the dyads and their coverage was calculated. Noise was removed using FFT filtering with parameter pcKeepComp=0.02 and peak detection was carried out with threshold 99%.

For each sample in an analysis set, sample-specific dyad positions obtained by nucleosome calling were enlarged to 20 bp and all positions were merged across the samples. Overlapping regions were joined. We excluded regions locating closer than 250 bp to tile borders and those residing in a region with high artifactual signals (chr III, 91000-93000 bp).

On this joint set of nucleosome dyads, we counted the number of overlapping fragments (reduced to their center position) for each sample. With x being the number of counts of sample-specific fragment centers overlapping one dyad region of the joint set and sum(x) being the sum of all counts across all dyad regions in the sample the data was normalized using the formula: normalized occupancy (dyad region) = log2(((x/sum(x))*1000)+0.001). The resulting matrix was subjected to principal component analysis. K-means clustering was applied to the resulting principal components to group nucleosomes based on similar occupancy patterns across sample conditions.

DNA shape features in windows of 320 bp around dyad positions were calculated with bioconductor/DNAshapeR (version 1.14.0). DNA rigidity scores of each position in windows of 320 bp around dyad positions were calculated as the length of the longest consecutive A*n*T*m* (*n*≥0, *m*≥0 and *n+m*≥2) sequence element that contains this position.

#### Data Availability

All raw and processed sequencing data generated in this study have been submitted to the NCBI Gene Expression Omnibus under accession numbers GSE145093 and GSE140614.

---

**Supplementary Figure 1.**
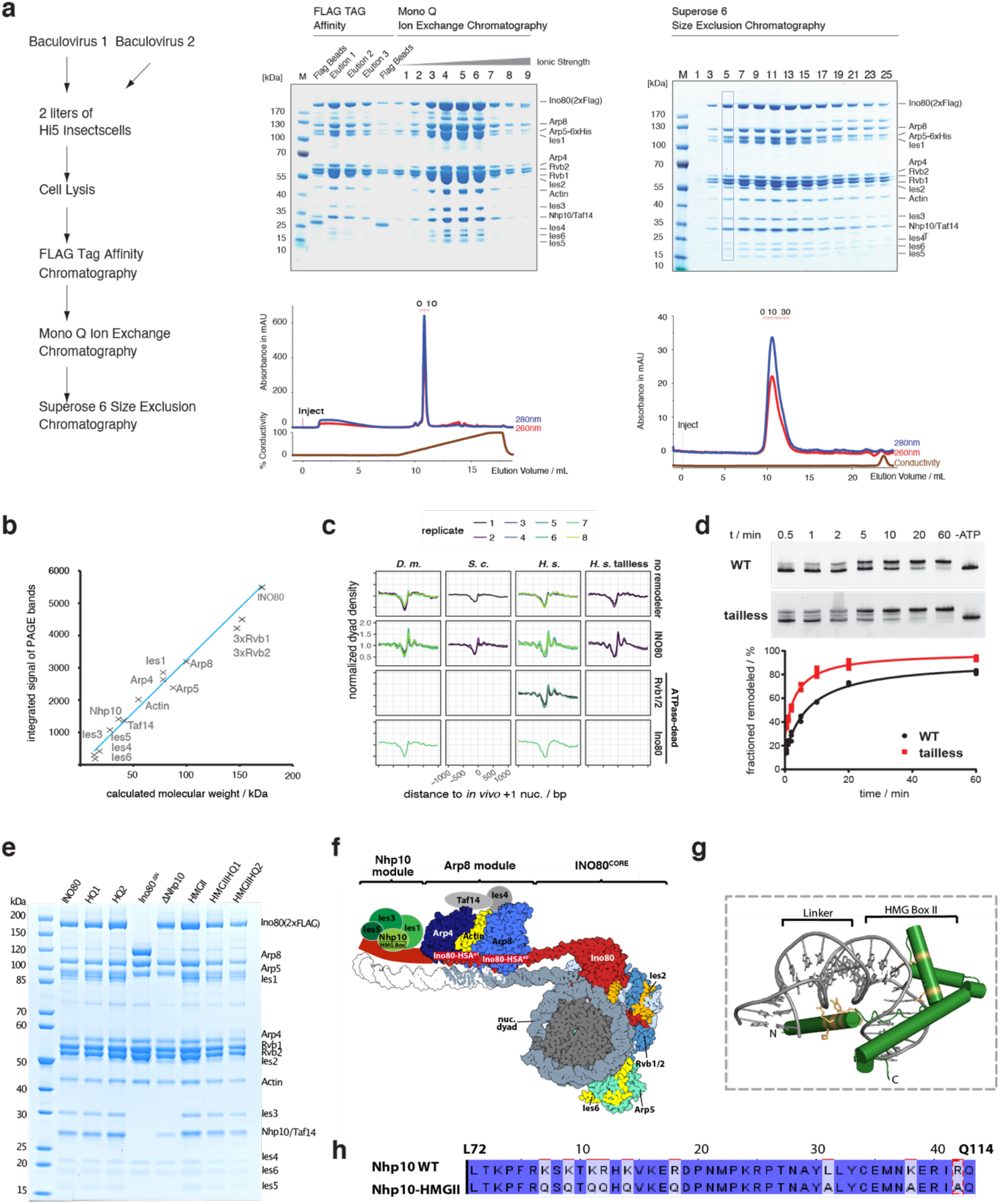
Expression, purification and activity of recombinant INO80. **a** Recombinant expression and purification of 15-subunit *S. cerevisiae* INO80 complex. Left: Schematic of expression and purification work flow. Two baculoviruses encoding five (Ino80, Arp5, Ies6, Rvb1 and Rvb2) and ten INO80 subunits (Ies1-5, Nhp10, Taf14, Actin, Arp4, Arp8), respectively, were used for insect cell expression. Middle and right: SDS-PAGE analysis of indicated chromatographies. Numbered lanes indicate elution fractions matching chromatograms below gels. Boxed lane represents a fraction used in this study. **b** Quantification of Coomassie-stained SDS PAGE bands shows stoichiometric assembly of recombinant *S.cerevisiae* INO80 complex. Note that AAA^+^ ATPase Rvb1 and Rvb2 form a hetero-hexamer. **c** Composite plots of MNase-seq data of individual replicates for the indicated combinations of histones (columns) and remodeling enzymes (rows). **d** top: Native gel electrophoresis analysis at indicated time points of mononucleosome sliding assay kinetics with wild type (WT) or tailless (tailless) recombinant *H. sapiens* histones and wild type recombinant *S. cerevisiae* INO80 complex. “-ATP” denotes 60 min time point without ATP. bottom: Quantification of data from top. **e** SDS-PAGE analysis of purified, recombinant WT (INO80) or indicated mutant complexes. **f** left: Structure-based ^6,16^ model of a nucleosome bound by the INO80 complex with indicated subunits. Taf14, Ies4 and Nhp10 module organization is assumed. **g** Model of Nhp10 HMG box-like and Linker region (residues 62-172) based on TFAM structure (pdb 3tq6). **h** Sequence alignment showing mutated residues in Nhp10-HMGII mutant. Panels e-h are also shown in the accompanying paper Oberbeckmann & Niebauer et al.

**Supplementary Figure 2.**
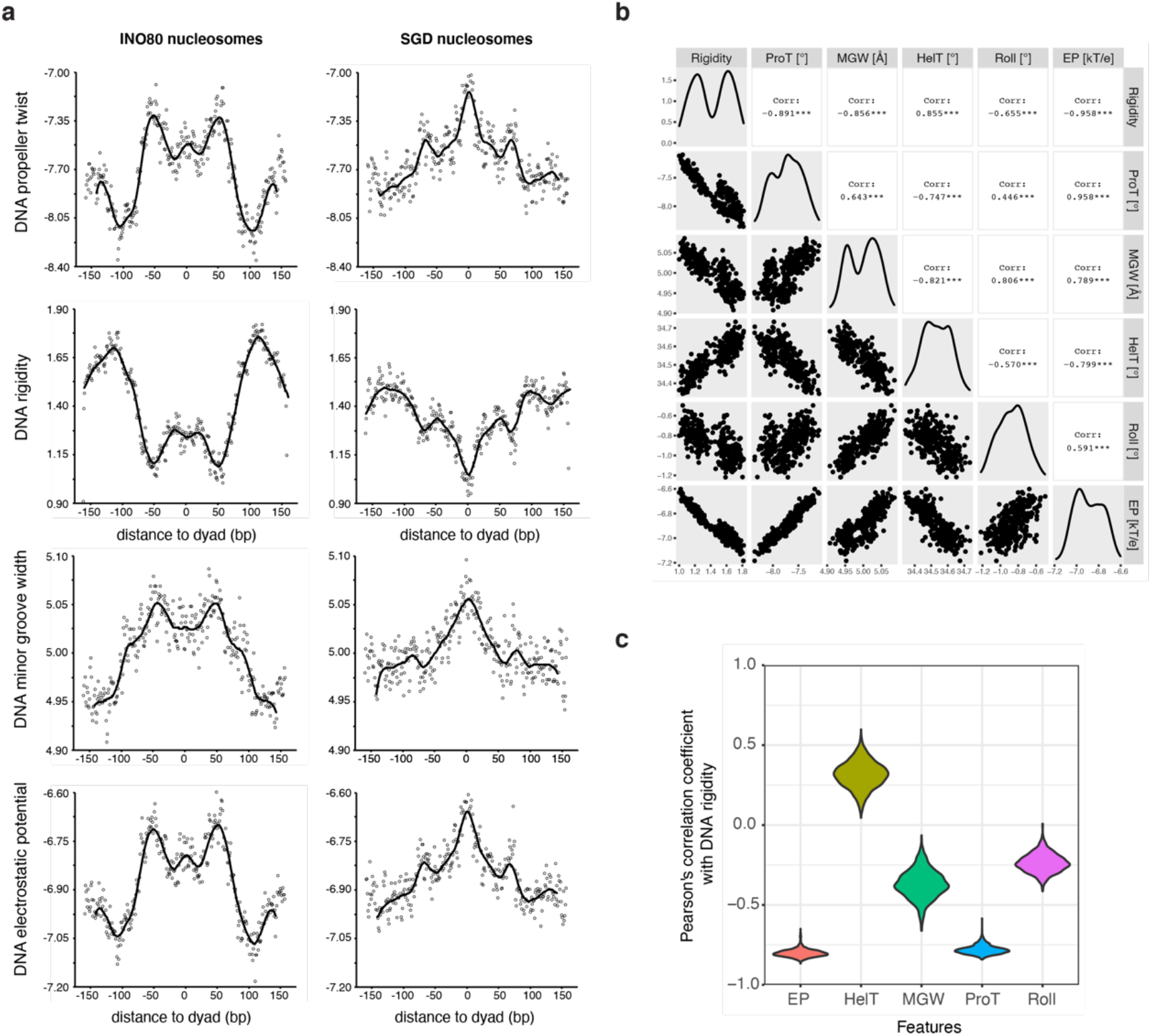
DNA shape/mechanics features of INO80 and SGD positioned nucleosomes. **a** DNA shape/mechanics profiles (DNA propeller twist, DNA rigidity, DNA minor groove width and DNA electrostatic potential) derived from INO80 and SGD positioned nucleosomes. **b** Pearson’s correlation coefficients between six DNA features: minor groove width (MGW), helix twist (HelT), propeller twist (ProT), Roll, Electrostatic potential (EP), and DNA rigidity. The average profiles of DNA features across all nucleosomal sequences are used to obtain the correlation coefficients between features. **c** Violin plot of Pearson’s correlation coefficients between DNA rigidity and other DNA features of all nucleosomal sequences. The coefficient is obtained by correlating the DNA feature profiles of each sequence individually.

**Supplementary Figure 3.**
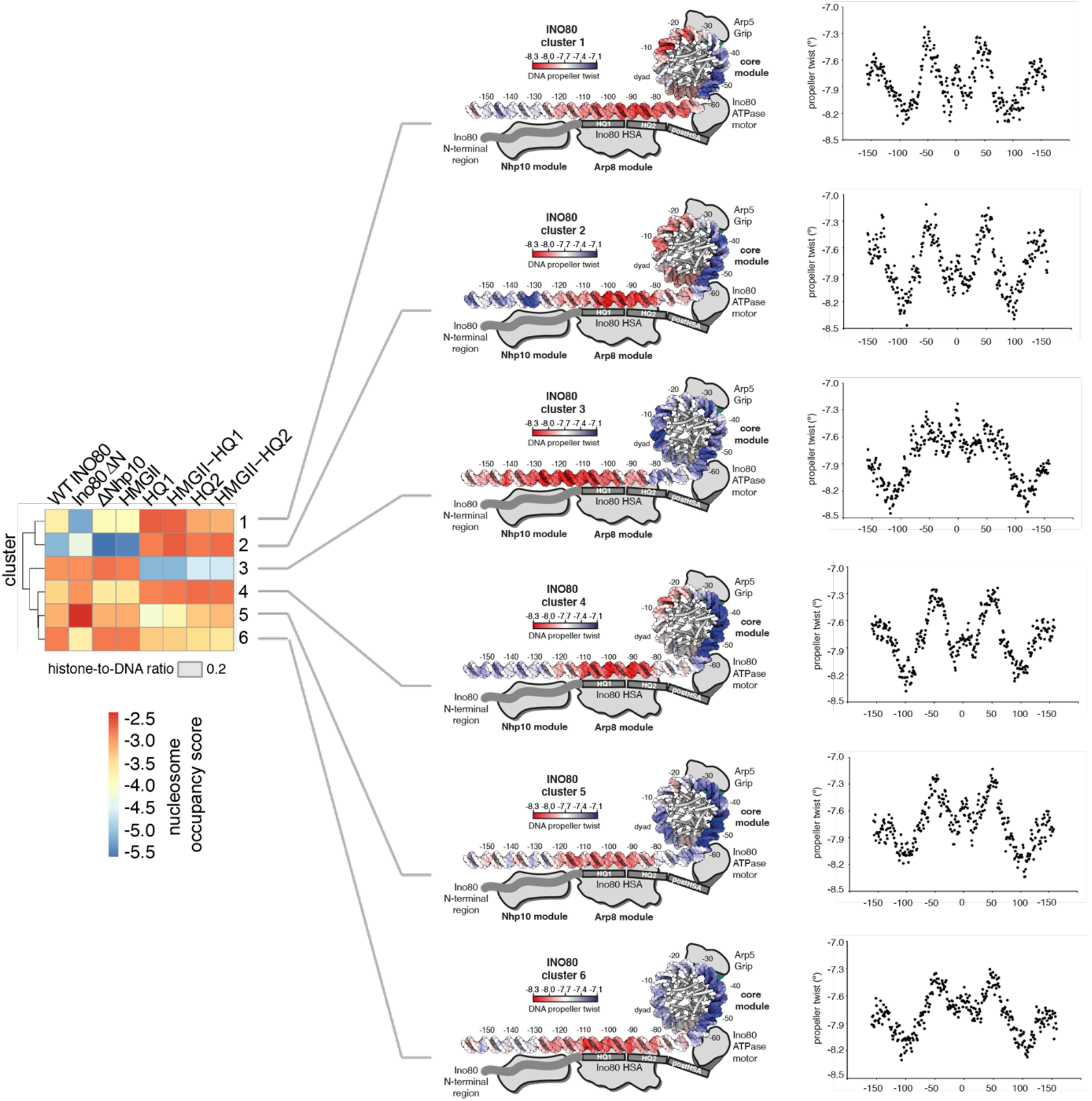
Clustering and DNA shape analysis of WT and mutant INO80. DNA propeller twist shape profile of nucleosomal DNA sequences. Color-coded mapping is shown for each cluster.

**Supplementary Figure 4.**
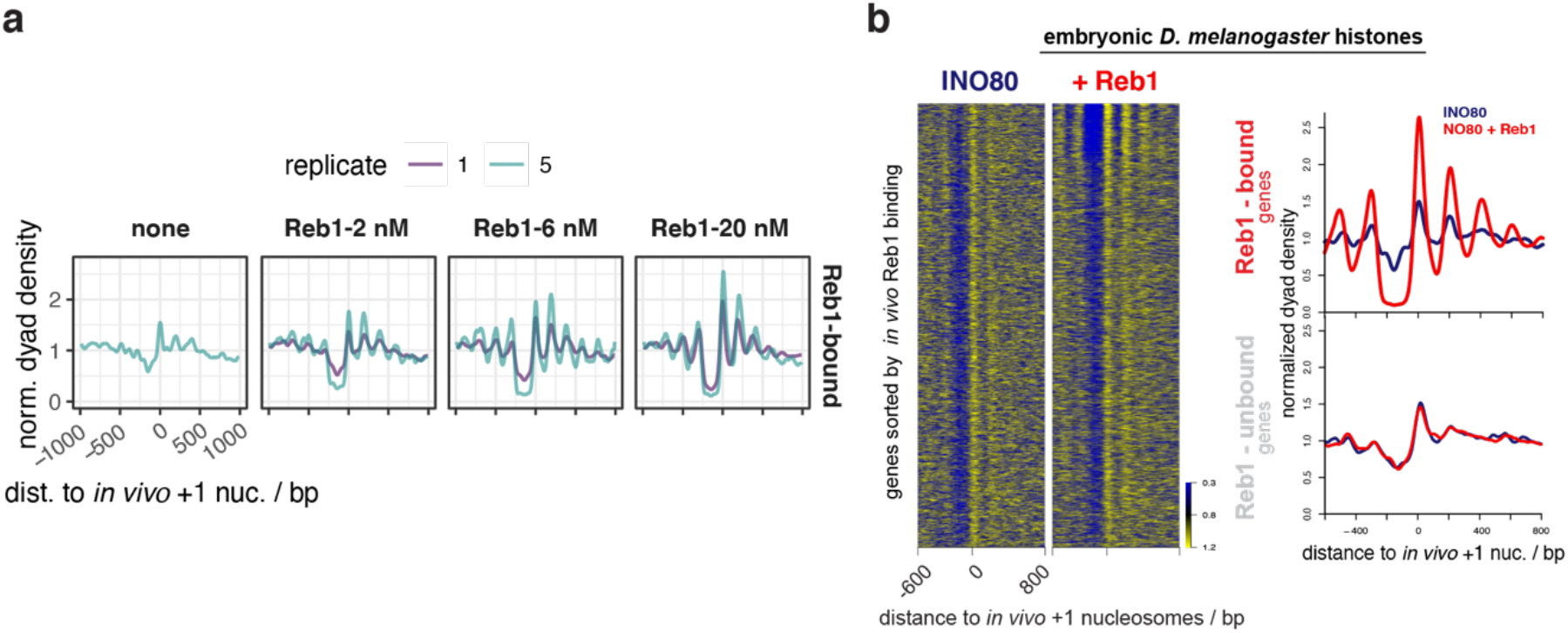
Nucleosome positioning in presence of Reb1. **a** Composite plots of MNase-seq data for individual replicates of samples as in Figure 6a,b, but only for genes with promoter Reb1 sites (Reb1-bound, same as red shading in Figure 6a) and also including SGD chromatin incubated with INO80 in the absence of Reb1 (none). **b** As Figure 6a,b but for the SGD chromatin with embryonic *D. melanogaster* histones at histone-to-DNA mass ratio of 0.4 and only 20 nM Reb1 (+ Reb1).

